# Reshaping the tumor microenvironment by degrading glycoimmune checkpoints Siglec-7 and -9

**DOI:** 10.1101/2024.10.11.617879

**Authors:** Chao Wang, Yingqin Hou, Jaroslav Zak, Qinheng Zheng, Kelli A. McCord, Mengyao Wu, Ding Zhang, Shereen Chung, Yujie Shi, Jinfeng Ye, Yunlong Zhao, Stephanie Hajjar, Ian A. Wilson, James C. Paulson, John R. Teijaro, Xu Zhou, K. Barry Sharpless, Matthew S. Macauley, Peng Wu

## Abstract

Cancer treatment has been rapidly transformed by the development of immune checkpoint inhibitors targeting CTLA-4 and PD-1/PD-L1. However, many patients fail to respond, especially those with an immunosuppressive tumor microenvironment (TME), suggesting the existence of additional immune checkpoints that act through orthogonal mechanisms. Sialic acid-binding immunoglobulin-like lectin (Siglec)-7 and -9 are newly designated glycoimmune checkpoints that are abundantly expressed by tumor-infiltrating myeloid cells. We discovered that T cells express only basal levels of Siglec transcripts; instead, they acquire Siglec-7 and -9 from interacting myeloid cells in the TME via trogocytosis, which impairs their activation and effector function. Mechanistically, Siglec-7 and -9 suppress T cell activity by dephosphorylating T cell receptor (TCR)-related signaling cascades. Using sulfur fluoride exchange (SuFEx) click chemistry, we developed a ligand that binds to Siglec-7 and -9 with high-affinity and exclusive specificity. Using this ligand, we constructed a Siglec-7/9 degrader that targets membrane Siglec-7 and -9 to the lysosome for degradation. Administration of this degrader induced efficient Siglec degradation in both T cells and myeloid cells in the TME. We found that Siglec-7/9 degradation has a negligible effect on macrophage phagocytosis, but significantly enhances T cell anti-tumor immunity. The degrader, particularly when combined with anti-CTLA-4, enhanced macrophage antigen presentation, reshaped the TME, and resulted in long-lasting T cell memory and excellent tumor control in multiple murine tumor models. These findings underscore the need to consider exogenous checkpoints acquired by T cells in the TME when selecting specific checkpoint blockade therapy to enhance T cell immunity.

## Main Text

Immune checkpoint blockade of CTLA-4 and PD-1/PD-L1 has demonstrated promising outcomes in diverse cancer patient populations^1^. However, response to these therapies is limited in individuals with immunosuppressive tumor microenvironments (TMEs), such as those notoriously found in glioblastoma and pancreatic tumors^2, 3^. One factor contributing to this resistance is the potential existence of additional immune checkpoints in the TME that operate independently of these well-established ones. Recently, the sialic acid-binding immunoglobulin-like lectin (Siglec) family members of glycan-binding proteins have been identified as glycoimmune checkpoints^4, 5^. Inhibitory Siglecs on immune cells bind to sialoglycans aberrantly expressed on tumor cells in a manner similar to PD-1/PD-L1 engagement, and trigger inhibitory signaling that suppress immune responses, contributing to immunosuppression^6, 7^. However, the impact of this inhibition on the crosstalk between innate and adaptive immunity in the cancer immunity cycle remains obscure.

## T cells acquire Siglec-7/9 receptors from myeloid cells

Examination of publicly available scRNA-seq datasets from pancreatic cancer patients^8^, revealed that *SIGLEC7* and *SIGLEC9* are primarily expressed in myeloid cells, particularly macrophages, with a detectable but very low expression in T cells^9^ (Fig. 1a,b). This pattern of expression also extends to other human solid tumors, including glioblastoma^10^, breast^11^, and colon^12^ cancers (Extended Data Fig. 1). However, to our surprise, by staining a pancreatic adenocarcinoma tissue array with anti-Siglec-7 and anti-Siglec-9 antibodies, we found high levels of both Siglecs presented on tumor-infiltrating T cells (Fig. 1c,d), suggesting the possibility that T cells may have acquired these Siglec receptors from surrounding Siglec-7/9^+^ myeloid cells through a process known as trogocytosis^13^.

**Fig. 1.**
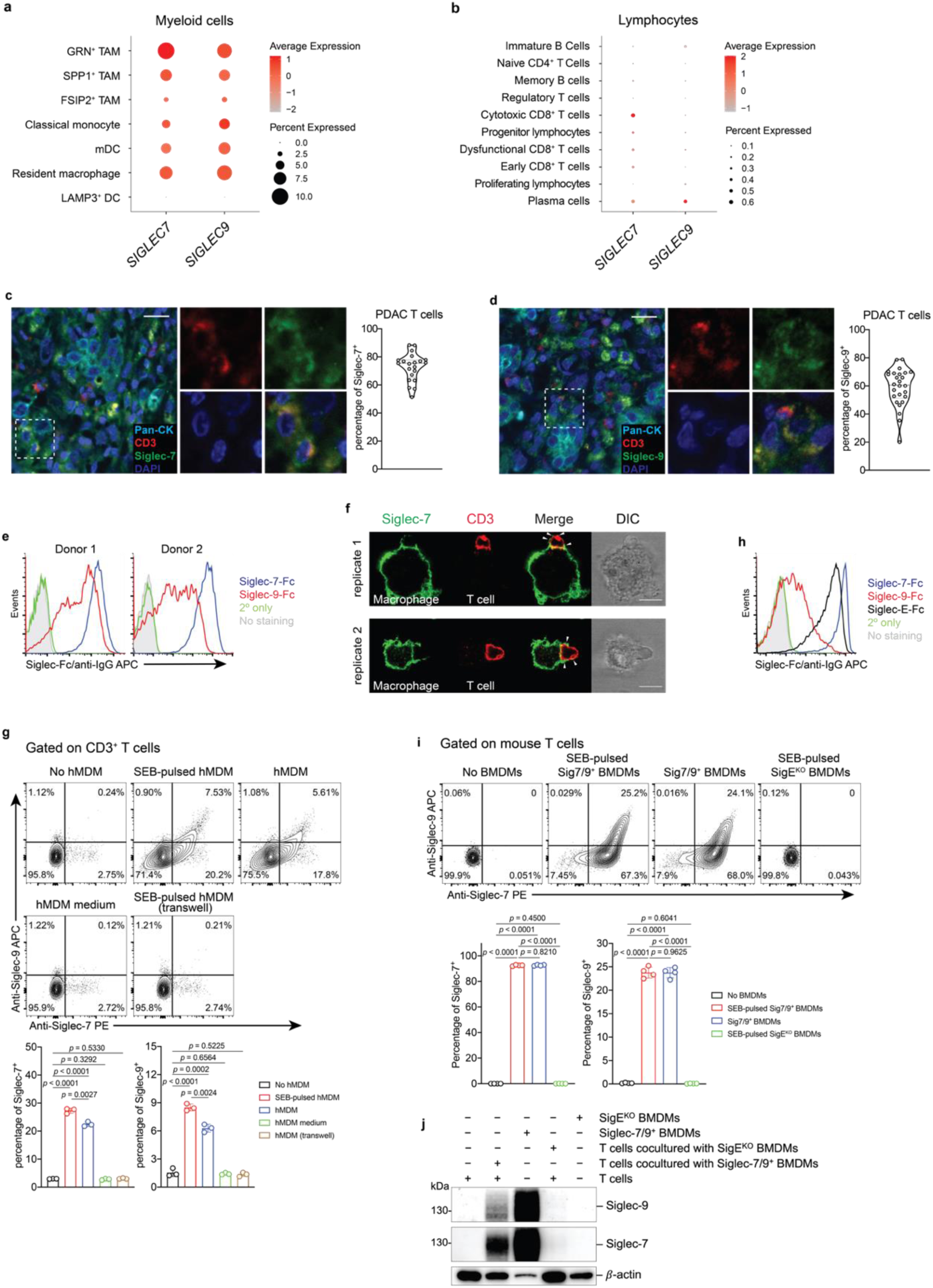
| Myeloid-associated Siglec-7/9 receptors are trogocytosed by neighboring T cells. **a**,**b**, Dot plots presenting the expression of *SIGLEC7 and SIGLEC9* genes across annotated tumor-infiltrating myeloid cells (**a**) and lymphocytes (**b**) from PDAC patients using 7 integrated datasets (PMID: 37633924). Dot radius is proportional to the percentage of each cell type expressing the *SIGLEC* gene, with average gene expression values depicted on the color gradient. **c**,**d**, Fluorescence microscopy imaging analysis of Siglec-7 and -9 presence on T cells from grade II and III PDAC tumor tissues. Scale bar, 20 µm. **e**, Analysis of cell-surface expression of Siglec-7 and -9 ligands on T cells from PBMCs of healthy donors by staining with Siglec-7Fc and Siglec-9Fc respectively. PBMC, peripheral blood mononuclear cell. **f**, Fluorescence microscopy imaging of Siglec-7 localization after 30 min coculture of SEB superantigen-pulsed hMDMs and donor-matched PBMC T cells that were pre-FACS sorted as Siglec-7/9^−^ population (there are 3∼4% Siglec-7^+^ T cells in the PBMCs of healthy donors). Scale bar, 10 µm. SEB, staphylococcal enterotoxin. **g**, Flow cytometry-based quantification of Siglec-7/-9 trogocytosis by T cells after 30 min of coculture with hMDMs (with or without SEB pulsing), or with medium isolated from hMDM culture, or with SEB-pulsed hMDMs separated by a transwell filter. **h**, Analysis of cell-surface expression of Siglec-7, Siglec-9 and Siglec-E ligands on WT mouse T cells by staining with Siglec-7Fc, Siglec-9Fc and Siglec-E-Fc, respectively. **i**, Flow cytometry analysis of Siglec-7/-9 trogocytosis by WT mouse T cells after 5 min of coculture with Sig7/9^+^ mouse BMDMs *vs.* SigE^KO^ BMDMs with or without SEB priming. BMDM, bone marrow-derived macrophage. **j**, Western blot analysis of Siglec-7 and -9 transferred to mouse T cells from Sig7/9^+^ BMDMs versus SigE^KO^ BMDMs. Data are mean ± s.d. Two-tailed unpaired Student’s *t*-test (**g**,**i**).

To investigate the possibility of Siglec-7/9 trogocytosis by T cells, we cocultured peripheral T cells that express abundant Siglec-7/9 ligands (Fig. 1e and Extended Data Fig. 2a) from healthy donors with staphylococcal enterotoxin B (SEB) superantigen-pulsed autologous human monocyte-derived macrophages (hMDMs) as the antigen-presenting cells (APCs). Siglec-7/9 molecules rapidly accumulated on interacting T cells, forming punctate structures rather than being uniformly distributed on the T cell surface (Fig. 1f and Extended Data Fig. 2b). Flow cytometry analysis also confirmed the rapid appearance of Siglec-7/9 on T cells, which occurred in the absence of antigen priming (Figure 1g and Extended Data Fig. 2c). By contrast, no Siglec-7/9 could be detected on T cells when incubated with collected hMDM culture medium or cocultured with hMDMs separated by transwell filters (Fig. 1g). Taken together, these observations provide compelling evidence that T cells acquire Siglec-7/9 molecules from neighboring macrophages via trogocytosis, which is dependent on direct cell-cell contact.

Similarly, when wild-type (WT) C57BL/6J (B6) mouse T cells that express high levels of Siglec-7/9 ligands (Figure 1h and Extended Data Fig. 2d), were cocultured with bone marrow-derived macrophages (BMDMs) or dendritic cells (BMDCs) from humanized Siglec-7/9 knock-in (Siglec-7^+^/-9^+^/Siglec-E knockout, hereafter referred to as Sig7/9^+^) B6 mice^14^, a large proportion of the T cells became Siglec-7 and -9 positive within five minutes (Fig. 1i and Extended Data Fig. 2e,f,g). Western blot analysis revealed that full-length Siglec-7 and -9 were acquired by T cells (Fig. 1j and Extended Data Fig. 2h,i). Consistent with what we had observed for human cells, Siglec-7/9 trogocytosis by T cells occurred to a similar extent regardless of antigen priming (Fig. 1i and Extended Data Fig. 2g), suggesting that this process is general, independent of antigen recognition, and primarily dependent on direct cell-cell engagement.

## *In vivo* Siglec-7/9 trogocytosis suppresses T cell effector function

To investigate if Siglec-7/9 trogocytosis by T cells also occurs *in vivo*, transgenic P14 T cells specific for the lymphocytic choriomeningitis virus (LCMV)-derived epitope gp33–41 (KAVYNFATC), were adoptively transferred into Sig7/9^+^ or control Siglec-E knockout (SigE^KO^) mice with established B16-GP33/GMCSF tumors (Fig. 2a). These tumors produce granulocyte-macrophage colony-stimulating factor (GMCSF), promoting the recruitment and differentiation of immunosuppressive TAMs^15^ as a source of Siglec-7/9 in the TME. Compared to P14 T cells isolated from SigE^KO^ mice, 2/3 of P14 T cells and 1/2 of the endogenous CD8^+^ T cells in the tumor-infiltrating lymphocytes (TILs) of Sig7/9^+^ mice were found to be Siglec-7/9^+^, respectively (Fig. 2b,c). In addition to TILs, both the adoptively transferred P14 and endogenous CD8^+^ T cells in the spleens and the tumor draining lymph nodes (dLNs) in Sig7/9^+^ mice were also found to be Siglec-7/9^+^ (Extended Data Fig. 3a,b,c).

**Fig. 2.**
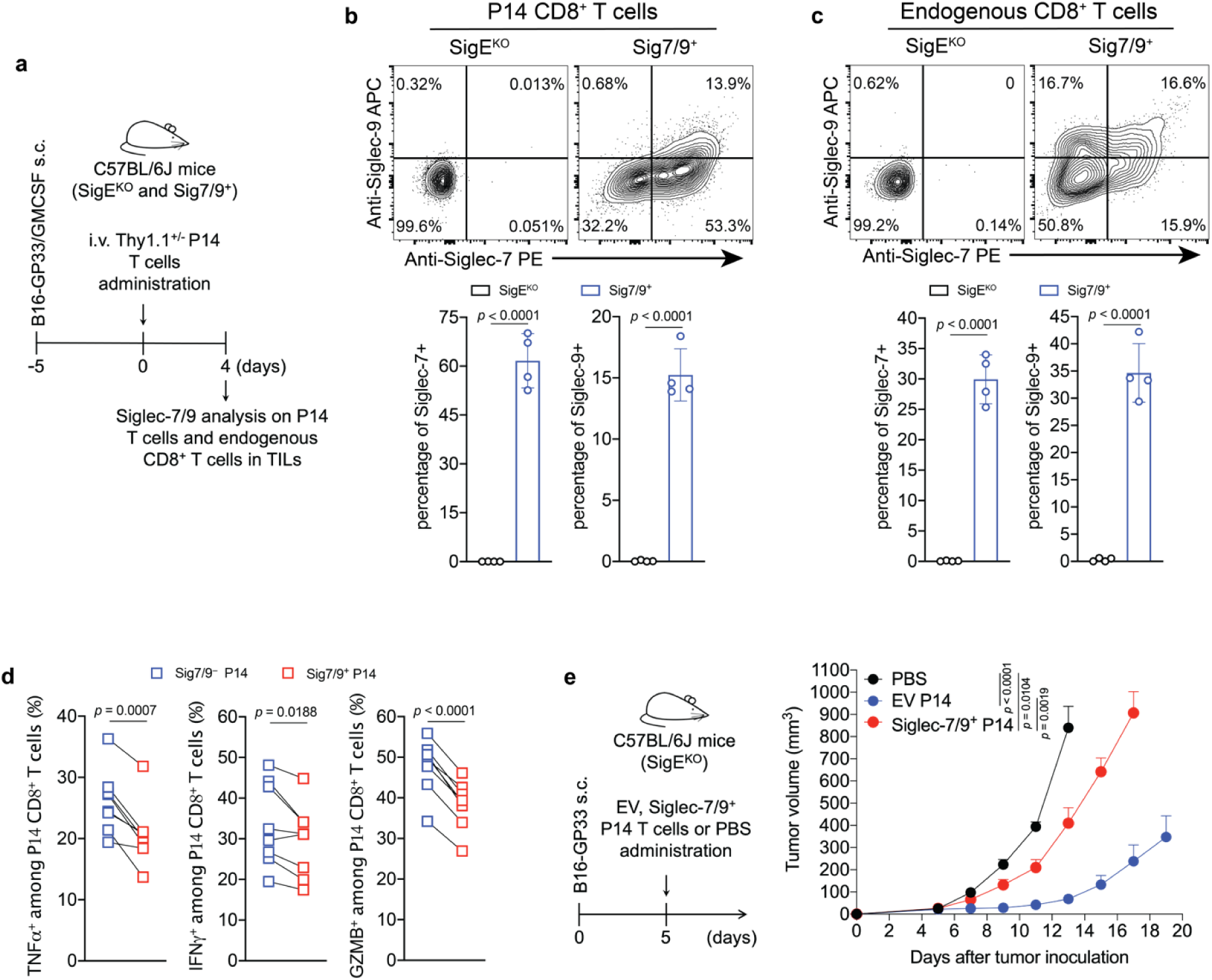
| *In vivo* Siglec-7/9 trogocytosis suppresses T cell effector functions. **a**, Experimental workflow of *in vivo* trogocytosis study. **b**, Analysis of Siglec-7/9 trogocytosis by adoptively transferred P14 CD8^+^ T cells (Thy1.1) in the TILs of B16-GP33/B16-GMCSF (9:1) tumors that were inoculated in SigE^KO^ and Sig7/9^+^ mice. TIL, infiltrating T lymphocyte. **c**, Analysis of Siglec-7/9 on endogenous CD8^+^ T cells in the TILs. **d**, Paired analysis of cytokine (IFNγ, TNFα and GZMB) production by Siglec-7/9^−^ and Siglec-7/9^+^ P14 CD8^+^ TILs. Data are mean ± s.d. Two-tailed unpaired Student’s *t*-test (**b**,**c**). Paired Student’s *t*-test (**d**). **e**, Evaluation of Siglec-7/9 co-expression on P14 T cells in the control of B16-GP33 tumors in SigE^KO^ mice (*n*= 5 mice for PBS, *n*= 6 mice for EV and Siglec-7/9^+^ P14 T cells, respectively). EV, empty vector. Average sizes of primary tumors ± SEM are presented in cubic millimeters (mm^3^) (**e**). Statistical analysis was performed using one-way ANOVA with Dunnett’s multiple comparisons test (**e**).

As myeloid cells, but not T cells, express high levels of Siglec-7 and -9 transcripts in Sig7/9^+^ mice (Extended Data Fig. 3d), we further examined the stability of Siglec-7/9 on these endogenous T cells by first labeling them with CFSE, followed by adoptive transfer into SigE^KO^ mice. After 24 hours, almost all Siglec-7 and -9 molecules disappeared from these transferred T cells isolated from the spleens and inguinal LNs of the recipient mice (Extended Data Fig. 3e,f). Conversely, SigE^KO^ T cells, upon being adoptively transferred to Sig7/9^+^ mice, gained both Siglecs to a level similar to that of endogenous T cells (Extended Data Fig. 3g, h). These observations strongly suggest that T cells do not produce Siglec-7 and -9, but rather acquire them from neighboring Siglec-7/9^+^ myeloid cells, and that the acquired Siglecs have a short half-life on T cells after trogocytosis in a Siglec-free environment.

To investigate the impact of trogocytosed Siglec-7/9 on T cell function, intratumoral and tumor dLN-infiltrating Siglec-7/9^+^ and Siglec-7/9^−^ P14 T cells were restimulated *ex vivo* with the gp33 peptide. Compared to Siglec-7/9^−^ T cells, Siglec-7/9^+^ cells significantly decreased the production of TNFα, IFNγ, granzyme B (GZMB), and IL-2 (Fig. 2d and Extended Data Fig. 3i). Furthermore, adoptive transfer of P14 T cells that were retrovirus-transduced to express both Siglecs into B16-GP33 bearing SigE^KO^ mice notably lost the capability for tumor control (Fig. 2e and Extended Data Fig. 3j). Taken together, these observations strongly suggest that T cell-acquired Siglec-7/9 are involved not only in the suppression of T cell effector function and anti-tumor immunity in the TME, but also in the inhibition of T cell activation in the tumor dLN.

Surprisingly, we observed that Siglec-7 and -9 mediate ligand-independent signaling in T cells. Upon anti-CD3/anti-CD28 stimulation, the activation of Siglec-7/9 ligand-free Jurkat T cells (Extended Data Fig. 4a) transduced with Siglec-7 or -9 (Extended Data Fig. 4b,c), was suppressed, as evidenced by reduced expression of the T-cell activation marker CD69 (Extended Data Fig. 4d). However, to block such inhibitory signaling, there is currently a lack of non-agonist anti-Siglec-9 antibodies. Although an anti-Siglec-7 antibody partially restored CD69 expression in Jurkat cells, an anti-Siglec-9 functional blocking antibody further decreased CD69 expression due to its agonistic property (Extended Data Fig. 4d). There are currently no blocking antibodies for both Siglec-7 and Siglec-9, nor are there tools to inhibit Siglec trogocytosis. Therefore, we sought an alternative strategy to directly block Siglec-7/9-mediated immune suppression in the TME by inducing targeted Siglec degradation. This involves the development of a heterobifunctional molecule with one arm binding to Siglec-7 and -9 with exclusive selectivity and the other arm for lysosomal targeting and degradation.

## Discovery of high-affinity and specific Siglec-7/9 ligands via SuFEx click chemistry

Given the close homology and analogous expression patterns of Siglec-7 and -9^16^, we envisioned that high-affinity ligands could be developed to bind both Siglecs with exclusive specificity. And these ligands could be further converted into degraders to induce Siglec-7/9 degradation in the TME, with the potential to restore antitumor immunity. However, natural sialoglycans bind to Siglecs with millimolar (mM) affinity and lack the required specificity^17^. Prior studies by Paulson *et al.* showed that derivatization of *N*-acetylneuraminic acid (Neu5Ac), the predominant sialic acid found in humans, at the C-5 or C-9 position using first-generation click chemistry, copper-catalyzed azide–alkyne cycloaddition (CuAAC), can generate high-affinity Siglec ligands^18^. Yet, many of these ligands lack the desired specificity and binding avidity. For instance, a high-affinity Siglec-7 ligand developed using this approach^19^ exhibited cross-binding to other Siglecs such as Siglec-2, -9 and -10 (Extended Data Fig. 5a,b). To address this issue, we chose to use second-generation click chemistry, the sulfur (VI) fluoride exchange (SuFEx)^20^, to produce a Neu5Ac-C9 derivative library by reacting 9-amino-Neu5Ac-α2-6-Gal-β1-4-GlcNAc (LacNAc) or 9-amino-Neu5Ac-α2-3-LacNAc with SOF_4_-derived electrophilic iminosulfur oxydifluorides (Fig. 3a). Instead of a triazole linkage created by CuAAC, the SuFEx transformation creates a sulfamide bond that could provide both H-bond donors and acceptors for Siglec binding. Leveraging on a ‘cell-surface screening platform’ (Extended Data Fig. 5c,d,e), we screened a library of sulfamide-linked Neu5Ac-LacNAc derivatives containing various heterocyclic substituents, and identified that the benzothiazole-modified ligands (**8** and **9**) with an α2-6-linked sialoside displayed substantially enhanced binding affinity to both Siglec-7 and -9 compared to the natural α2-6-Neu5Ac-LacNAc (**6**); whereas other derivatized ligands (**10**-**18**) showed varying degrees of comparatively low to moderate binding affinity (Fig. 3b,c and Extended Data Fig. 5f,g. The chemical synthesis of ligands is shown in the Supplemental Information).

**Fig. 3.**
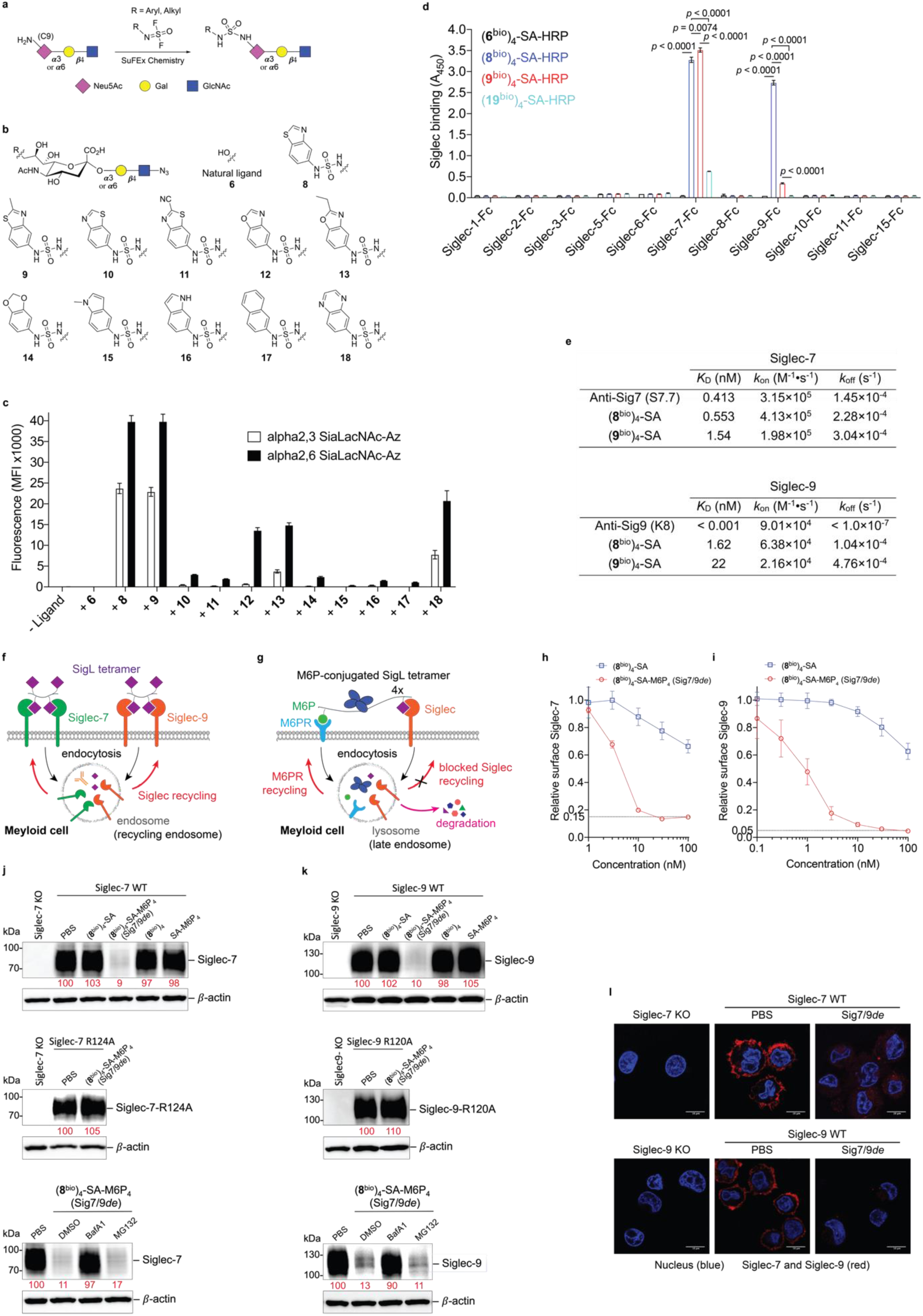
| Development of a Siglec-7/9 degrader via the discovery of high-affinity and selective Siglec-7/9 ligands. **a**, The synthesis of sulfamide linked α2-6 or α2-3 9-amino-Neu5Ac-LacNAc (9-amino-Neu5Ac-Galβ1-4GlcNAc) derivatives using SuFEx click chemistry, in which 9-amine tagged Neu5Ac is subjected to reaction with a library of iminosulfur oxydifluorides to form diversified sulfamide-linked Neu5Ac mimetics. SuFEx, Sulfur (VI) fluoride exchange; Neu5Ac, *N*-acetylneuraminic acid; Gal, galactose; GlcNAc, *N*-acetylglucosamine. **b**, The structures of a library of sulfamide-linked Neu5Ac-LacNAc derivatives. **c**, Measurement of Siglec-7 binding affinity of the synthetic Neu5Ac ligands (shown in **b**) installed on the cell surface by staining with Siglec-7Fc determined by MFI using flow cytometry. MFI, mean fluorescence intensity. **d**, An ELISA assay for Siglec cross-binding assessment of HRP-functionalized SigL tetramers. ELISA, enzyme-linked immunosorbent assay. Data are mean ± s.d. Two-tailed unpaired Student’s *t*-test. *ns*, not significant (**d**). **e**, Biolayer interferometry (BLI) assay for determination of the binding affinity *K*_D_ affiliated with rate constants of *k*_on_ (association), *k*_off_ (dissociation) for (**8^bio^**)_4_-SA and (**9^bio^**)_4_-SA tetramers and anti-Siglec-7/-9 antibodies towards Siglec-7 and Siglec-9, respectively. **f**, Schematic presentation of rapid internalization and recycling of Siglec-7/-9 in myeloid cells upon treatment with (SigL^bio^)_4_-SA tetramer. **g**, Schematic presentation of the rationale for degrading Siglec-7/-9 through the incorporation of M6P into the (SigL^bio^)_4_-SA tetramer, which targets and delivers Siglec-7/-9 to the lysosomes for degradation upon recognition by the M6P receptor (M6PR). **h**, Analysis of cell-surface Siglec-7 levels in Siglec-7^+^ U937-derived macrophages after treatment with (**8**^bio^)_4_-SA with or without M6P conjugation for 1 hour at various doses. **i**, Analysis of cell-surface Siglec-9 levels in Siglec-9^+^ U937-derived macrophages after treatment with (**8**^bio^)_4_-SA with or without M6P conjugation for 1 hour at various doses. **j**,**k**, West blot analysis of Siglec-7 and -9 in Siglec-7/-9 KO, WT and R-mutant U937-derived macrophages after treatment with 30 nM (**8**^bio^)_4_-SA-M6P_4_ (Sig7/9*de*) for 24 hours in comparison with ctrs (treated with PBS, (**8**^bio^)_4_-SA or SA-M6P_4_, and Siglec-KO cells), as wells as in the presence of lysosome inhibitor BafA1 or proteasome inhibitor MG132. **l**, Microscopic confirmation of the degradation of Siglec-7 and -9 in Siglec-expressing U937-derived macrophages, respectively. Scale bar, 10 µm.

## Siglec-7/9 ligand-assembled tetramers act as antibody surrogates for targeting Siglecs

Although natural glycoside monomers bind to their cell-surface receptors with low affinity, high affinity and specificity can be achieved through multivalent interactions^21^. Inspired by nature, we assembled glyco-tetramers using streptavidin (SA) and monomeric biotinylated Siglec ligand (SigL^bio^) (Extended Data Fig. 6a). Using the horseradish peroxidase (HRP)-conjugated (SigL^bio^)_4_-SA, we evaluated the cross-reactivity of the tetramers assembled from ligands **8** and **9** against a panel of human Siglec-Fc fusion proteins (Extended Data Fig. 6b,c and Fig. 3d). Both (**8**^bio^)_4_-SA and (**9**^bio^)_4_-SA tetramers exhibited exclusive binding specificity for Siglec-7 and -9, with (**8**^bio^)_4_-SA showing similar avidity for both Siglecs, whereas (**9**^bio^)_4_-SA showed higher avidity for Siglec-7 than Siglec-9 (Fig. 3d). By contrast, the tetramer assembled from the natural sialoside **6** lacked detectable binding to all Siglec-Fcs tested. And compared to (**8**^bio^)_4_-SA and (**9**^bio^)_4_-SA, the tetramer assembled from the only high-affinity Siglec-7 ligand **19**^19^ known to date (Extended Data Fig. 6d), showed significantly weaker binding to Siglec-7 (Fig. 3d and Extended Data Fig. 6c).

The binding avidities of (**8**^bio^)_4_-SA and (**9**^bio^)_4_-SA tetramers to Siglec-7/9 were found to be in the nanomolar (nM) range, comparable to that of the commercial anti-Siglec-7 and anti-Siglec-9 antibodies (Fig. 3e and Extended Data Fig. 6e). Both tetramers, when fluorescently tagged, enabled the detection of Siglec-7/9 on U937 cells in a manner similar to that of anti-Siglec antibodies (Extended Data Fig. 7a,b), whereas the (**19**^bio^)_4_-SA tetramer showed only weak labeling of Siglec-7^+^ cells (Extended Data Fig. 7a). In contrast to Siglec antibodies, which retained similar labeling ability to U937 cells expressing Siglec-7/9 mutants, wherein a key conserved arginine in the Siglec V-set binding domain is mutated to alanine, the (**8**^bio^)_4_-SA and (**9**^bio^)_4_-SA tetramers showed no binding (Extended Data Fig. 7a,b). This observation indicates that the binding of cell-surface Siglec by the tetramers relies on the canonical arginine-based salt bridge, which is essential for sialic acid recognition. Consistent with the binding data (Fig. 3d), (**8**^bio^)_4_-SA and (**9**^bio^)_4_-SA tetramers labeled Siglec-7/9^+^ cells with exclusive selectivity (Extended Data Fig. 7c,d,e). Therefore, (**8**^bio^)_4_-SA that binds to both Siglec-7 and -9 with similar affinity was chosen for the follow-up studies.

## Mannose-6-phosphate-functionalized SigL tetramers induce Siglec-7/9 degradation

We next investigated the fate of cell-surface Siglec-7 and -9 upon (**8**^bio^)_4_-SA-induced oligomerization. Incubating U937-derived macrophages expressing Siglec-7 and -9 with (**8**^bio^)_4_-SA-AF488 induced a rapid decrease of Siglec-7 and Siglec-9 from the cell surface (Extended Data Fig. 8a,b). Fluorescence microscopy imaging revealed internalization of AF488-associated fluorescence and Siglec molecules, colocalizing with the early endosome marker Rab5 (Extended Data Fig. 8c,d). After removal of the unbound tetramers, cell-surface expression of Siglec-7 and - 9 recovered within a few hours (Extended Data Fig. 8e,f,g,h), presumably via the recycling from early endosomes^22, 23^ (Fig. 3f).

To prevent recycling of the internalized Siglec-7/9, we developed a Siglec-7/9 degrader (Sig7/9*de*) based on the recently reported lysosome-targeting chimera (LYTAC) technology^24^. Sig7/9*de* was constructed by incorporating mannose-6-phosphate (M6P) onto the (**8**^bio^)_4_-SA tetramer to form (**8**^bio^)_4_-SA-M6P_4_ and direct the internalized Siglec-7/9 to the lysosome for degradation (Fig. 3g. Synthesis and characterization of SA-M6P_4_ are shown in the Supplemental Information). The M6P-mediated degradation relies on the shuttling of lysosome-associated cation-independent M6P receptor (CI-M6PR) between the cell surface and late endosomes^25^ (Extended Data Fig. 9a). Thus, the suppressed functions of myeloid cells will be unleashed particularly in the TME with upregulated sialoglycans. We treated Siglec-7^+^ and -9^+^ U937-derived macrophages with Sig7/9*de* and observed nearly complete depletion of cell-surface Siglec-7/9 molecules within 1 hour in a dose-dependent manner. By contrast, the control tetramer lacking M6P removed only a small fraction of Siglecs from the cell surface (Fig. 3h,i). Western blot analysis confirmed near-complete degradation of Siglec-7/9 in Sig7/9*de*-treated macrophages compared to the control groups (treated with PBS, (**8**^bio^)_4_-SA or SA-M6P_4_, and Siglec-KO cells) (Fig. 3j,k). The degradation occurred in less than 4 hours with maximum degradation achieved upon a 24-hour treatment (Extended Data Fig. 9b). The Siglec-7/9-R mutants with disrupted Siglec-sialic acid interactions were resistant to Sig7/9*de*-induced degradation (Fig. 3j,k). And the degradation was blocked by the lysosome inhibitor bafilomycin A1 (BafA1), but not the proteasome inhibitor MG132 (Fig. 3j,k), consistent with a mechanism involving lysosome-mediated degradation^24^. Microscopic examination further confirmed Siglec-7/9 degradation as opposed to endocytosis (Fig. 3l). After removal of the degrader, recovery of Siglec-7/9 occurred gradually over 3 days (Extended Data Fig. 9c), allowing follow-up studies on mitigating their inhibitory effects to be examined within a defined timeframe.

## Siglec-7/9 degradation exhibits limited impact on macrophage phagocytosis of cancer cells

Macrophages have been increasingly explored as potential candidates for cancer immunotherapy due to their ability to recognize and eliminate transformed cells via phagocytosis^26,27^. However, cancer cells aberrantly express sialoglycans as a “don’t eat me” signal, preventing the attack by immune cells including macrophages^4, 28^. These sialoglycans either shield tumor cells like a protective barrier^29^ or engage with inhibitory Siglec receptors to suppress immune cell function^7, 30, 31^. Despite previous studies, the direct role of Siglec-7/9 in macrophage-mediated anti-tumor immunity remains obscure. To directly investigate whether Siglec-7/9 degradation is capable of facilitating macrophage phagocytosis of cancer cells expressing Siglec-7/9 ligands (Siglec-7/9Ls), we treated hMDMs with Sig7/9*de*, which resulted in efficient degradation of both Siglec-7 and -9 simultaneously (Fig. 4a,b,c). Microscopy and flow cytometry-based phagocytosis assays were utilized to evaluate the phagocytic capacity against a panel of Siglec-7/9L^+^ cancer cells, including colon cancer, breast cancer, pancreatic cancer, glioblastoma, ovarian cancer, B-lymphoma and T-cell acute lymphoblastic leukemia cell lines (Extended Data Fig. 10a,b,c,d). Unexpectedly, we found that Siglec-7/9 degradation was not sufficient to induce phagocytosis in all cancer cell lines tested (Extended Data Fig. 10d,e), suggesting a minor role for Siglec-7/9 in macrophage-mediated tumor phagocytosis.

**Fig. 4.**
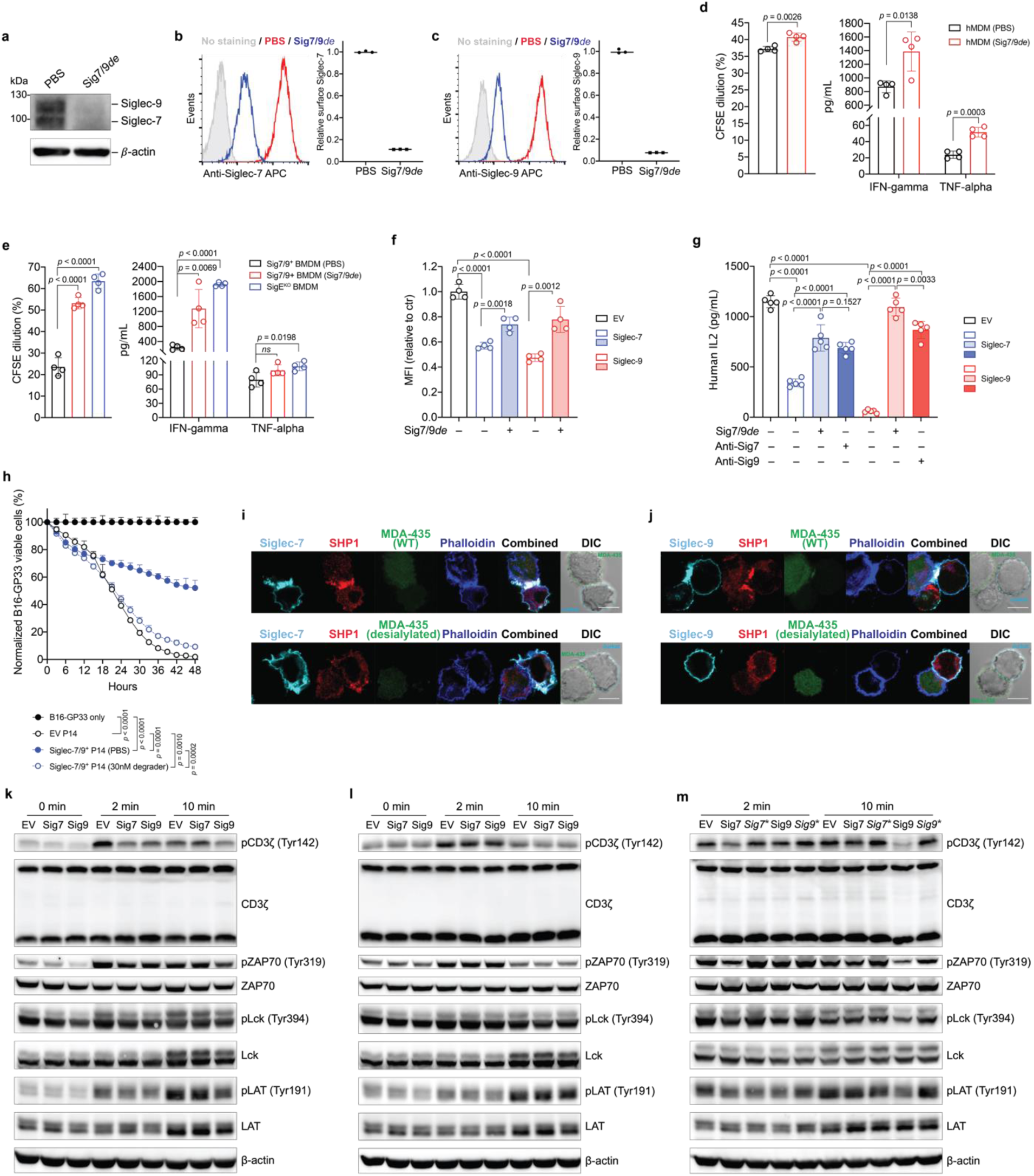
| Siglec-7/9 degradation rescues impaired T cell functions caused by Siglec trogocytosis. **a**, Western blot analysis of Siglec-7/-9 in human monocyte-derived macrophages (hMDMs) upon treatment with 30 nM Sig7/9*de* for 24 hours. **b**,**c**, Analysis of cell-surface Siglec-7/-9 levels on hMDMs upon treatment with 30 nM Sig7/9*de* for 1 hour. **d**, Human CD8^+^ T cell activation (proliferation and effector cytokine production) by anti-CD3 (OKT3) stimulation in coculture with donor-matched hMDMs in the absence or presence of 30 nM Sig7/9*de* for 72 hours. **e**, P14 CD8^+^ T cell activation (proliferation and cytokine production) by GP33 peptide (KAVYNFATM) stimulation in coculture with Sig7/9^+^ mouse BMDMs in the absence or presence of 30 nM Sig7/9*de* for 48 hours compared to coculture with SigE^KO^ BMDMs. **f**, Evaluation of Jurkat T cell (EV, Siglec-7 WT and Siglec-9 WT) activation, indicated by CD69 expression, upon OKT3/anti-CD28 stimulation for 24 hours in the absence or presence of 30 nM Sig7/9*de*. **g**, Assessment of Siglec degradation (Sig7/9*de* treatment at 30 nM) or blockage (anti-Sig7 IE8 or anti-Sig9 mAbA treatment at 30 nM) in restoration of IL2 secretion by Jurkat T cells cocultured with MDA-MB-435 cells (HER2^+^) for 24 hours in the presence of anti-HER2/anti-CD3 bispecific T cell engager (anti-HER2 BiTE) and anti-CD28. **h**, Assessment of Siglec-7/9^+^ P14 T cell-mediated killing of B16-GP33 tumor cells (E:T ratio = 5:1) in the presence or absence of 30 nM Sig7/9*de*. Data are mean ± s.d. Two-tailed unpaired Student’s *t*-test (**d**,**e**,**f**,**g**,**h**). **i**,**j**, Fluorescence microscopic imaging of Siglec-7 (**i**) and Siglec-9 (**j**) localization as well as SHP1 recruitment at the immunological synapse between Siglec-expressing Jurkat cells and MDA-MB-435 (HER2^+^) cancer cells after coculture at 37 °C for 30 min in the presence of anti-HER2 BiTE and anti-CD28 . Scale bar, 10 μm. SHP, Src homology 2 containing protein tyrosine phosphatase. **k**,**l**,**m**, Immunoblot analysis of the phosphorylation status of TCR signaling components (CD3ζ, ZAP70, Lck and LAT) in Jurkat T cells (EV, Sig7 and Sig9) cocultured with HER2^+^ MDA-MB-435 cancer cells in the presence of anti-HER2 BiTE and anti-CD28 at various time points, in which each experimental condition consisted of an equal number of Jurkat and cancer cells at a 1:1 ratio. Jurkat (EV, Sig7, Sig9) coculture with WT Siglec-7/9L^+^ MDA-MB-435 cells (**k**), Jurkat (EV, Sig7, Sig9) coculture with pre-desialylated MDA-MB-435 cells (**l**), Jurkat (EV, Sig7, *Sig7*\*, Sig9, *Sig9*\*) coculture with WT Siglec-7/9L^+^ MDA-MB-435 cells, in which *Sig7*\* and *Sig9*\* indicate Siglec-7 and Siglec-9 pre-degradation respectively (**m**).

## Siglec-7/9 degradation restores T cell activation and effector function

Next, we investigated the impact of degradation of myeloid-associated Siglec-7/9 on T cell function. Coculturing human peripheral CD8^+^ T cells with autologous hMDMs treated with Sig7/9*de* not only significantly increased the expression of T cell activation makers, CD69, CD25, and PD-1 (Extended Data Fig. 11a,b,c), but also led to enhanced T cell proliferation and cytokine production, e.g., IFNγ and TNFα (Fig. 4d). Similarly, treating cocultured P14 T cells and gp33 peptide-pulsed BMDMs from Sig7/9^+^ mice with Sig7/9*de* (Extended Data Fig. 12a,b) also increased T cell proliferation and IFNγ secretion to levels comparable to those observed when T cells were cocultured with SigE^KO^ BMDMs (Fig. 4e).

The above experiment, however, did not provide information on the direct impact of Siglec-7/9 degradation on T cells. To address this question, Jurkat T cells transduced with Siglec-7 or -9 were treated with Sig7/9*de*, which largely restored CD69 upregulation following anti-CD3/anti-CD28 stimulation (Extended Data Fig. 11d,e and Fig. 4f). Engagement with *trans* sialoglycan ligands is the primary trigger of Siglec-7 and -9-mediated inhibitory signaling. When cocultured with HER2^+^ MDA-MB-435 cancer cells expressing high levels of Siglec-7/9Ls (Extended Data Fig. 11f) in the presence of bispecific anti-HER2/anti-CD3 T cell engager (anti-HER2 BiTE)^29^ and anti-CD28, Siglec-7^+^ or -9^+^ Jurkat cells significantly reduced IL-2 production compared to their empty vector (EV)-transduced counterparts (Fig. 4g). Arming anti-HER2 BiTE with a sialidase^29^, which primarily removes sialylated glycans on the cancer cells, largely restored IL-2 production (Extended Data Fig. 11g,h). Likewise, the addition of Sig7/9*de* also restored Jurkat IL-2 production, and the restoration was greater than that achieved with Siglec blocking antibodies, especially when anti-Siglec-9 was used (Fig. 4g). Similarly, Siglec-7/-9 co-expression in P14 T cells led to dampened cytotoxicity against Siglec-7/9L^+^ B16-GP33 tumor cells (Extended Data Fig. 12c and Fig. 4h), which can be significantly rescued by Sig7/9*de* treatment (Extended Data Fig. 12d,e,f and Fig. 4h).

To further investigate the molecular mechanism by which Siglec-7/9 degradation rescues T cell function, we performed microscopic imaging studies, which revealed that Siglec-7 and -9 accumulate primarily at the immunological synapse between Jurkat cells and cancer cells, accompanied by the recruitment of the phosphatase SHP-1 to the synapse (Fig. 4i,j). Recruitment of SHP phosphatases to the immunological synapse by PD-1 is known to be responsible for TCR dephosphorylation, thereby suppressing its downstream signaling^32^. This is also the case for recruitment of SHP-1 to the synapse by Siglec-7 and -9, with Siglec-9 showing a more pronounced inhibitory effect (Fig. 4g). After 2- and 10-minute coculturing Siglec-7^+^ or -9^+^ and Siglec^−^ Jurkat cells with MDA-MB-435 cells in the presence of anti-HER2 BiTE and anti-CD28, it was found that the phosphorylation of CD3ζ and the proximal TCR signaling components ZAP-70, Lck, and LAT was considerably lower in Siglec^+^ cells (Fig. 4k), aligning with the observed inhibition of T cell activation and cytokine production (Fig. 4d,g). Desialylating target cells is known to facilitate T cell effector functions^29^. Pre-desialylation of MDA-MB-435 cells largely relieved the reduced phosphorylation of CD3ζ and ZAP-70, but failed to fully restore the phosphorylation of Lck (Fig. 4l). Intriguingly, Siglec-7 or -9 pre-degradation on Jurkat T cells effectively restored the phosphorylation of all TCR signaling components examined (Fig. 4k), suggesting that Siglec degradation may be a more efficient way for restoring T cell function.

## *In vivo* Siglec-7/9 degradation in TME

After confirming that Siglec-7/9 degradation is able to enhance T cell activation and effector function *in vitro*, we investigated whether Siglec-7/9 degradation could be accomplished *in vivo*. Siglec-7/9L tetramer ((**8**^bio^)_4_-SA) showed no cross-binding to mouse Siglec receptors, thus excluding any off-target effects (Fig. 5a and Extended Data Fig. 13a). We then administered Sig7/9*de* intratumorally in B16-GMCSF tumors established in Sig7/9^+^ mice, which are characterized by an abundant presence of myeloid cells^15^ (Fig. 5b,c and Extended Data Fig. 13b), in particular, Siglec-7/9^+^ TAMs (Extended Data Fig. 13b and Fig. 5d), mimicking the microenvironment commonly found in many human tumor types^33^. Maximum Siglec-7/9 degradation, reflected by cell-surface reduction of approximately 60% Siglec-7 and 70% Siglec-9, was achieved at a low dose (10 μg per tumor) on day 2 following the degrader administration, after which the newly synthesized Siglec-7/9 began to repopulation (Fig. 5c). Within CD11b^+^ cells, notable Siglec-7/9 degradation was observed across multiple subsets, including TAMs, MDSCs, and DCs (Fig. 5d). Additionally, significant Siglec-7/9 degradation was also observed in tumor infiltrating T cells and T cells in the tumor dLNs (Fig. 5e,f), as well as in natural killer (NK) cells although to a lesser extent (Extended Data Fig. 13c,d).

**Fig. 5.**
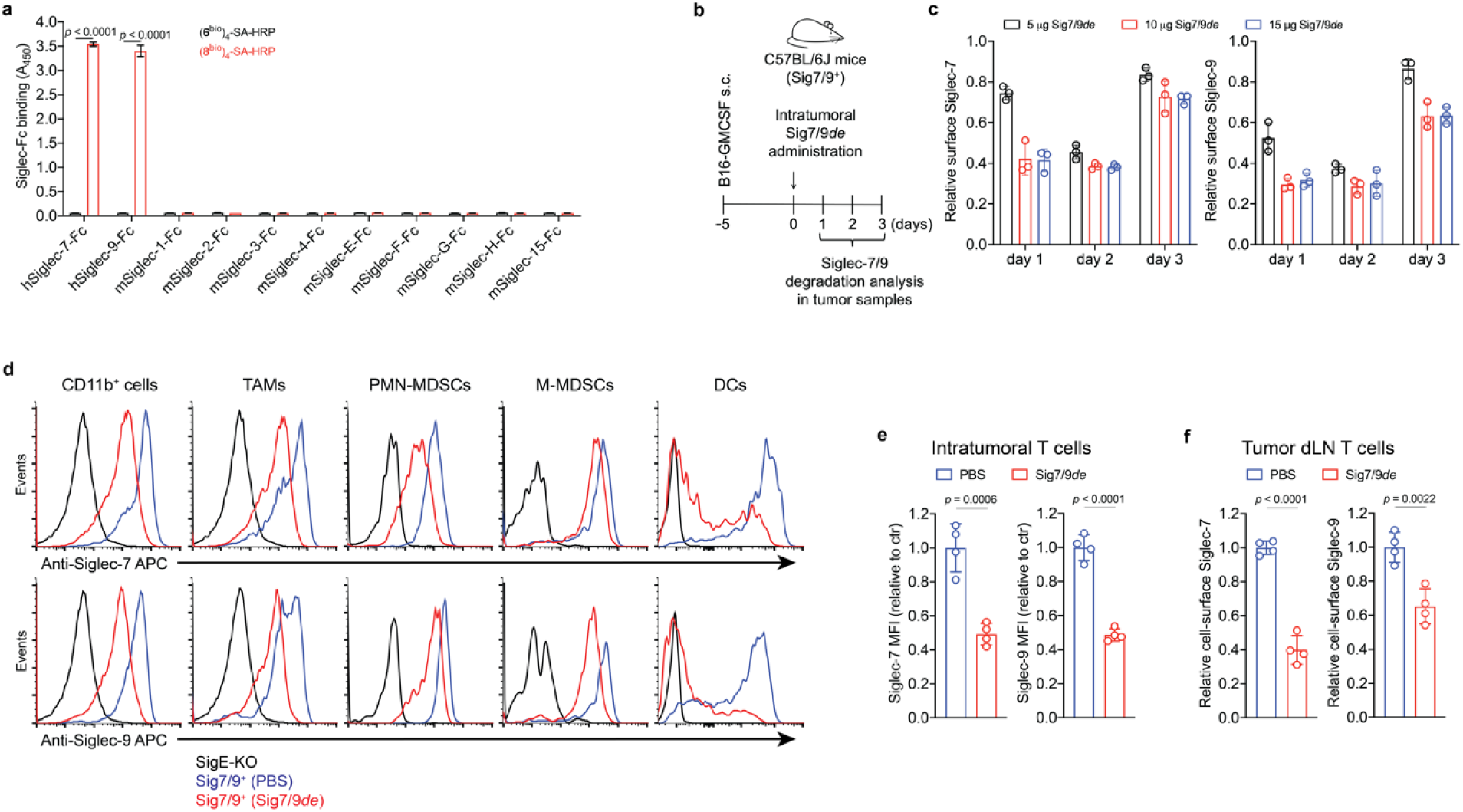
| Sig7/9*de* induces efficient *in vivo* Siglec-7/9 degradation. **a**, An ELISA assay for analyzing the cross-binding of tetramers assembled from the high-affinity Siglec-7/9 ligand (**8**^bio^) or the natural Neu5Ac-LacNAc (**6**^bio^) towards a panel of mouse Siglecs and human Siglec-7 and - 9. **b**,**c**, Analysis of *in vivo* Siglec-7/-9 degradation efficiency in tumor-infiltrating CD11b^+^ cells from B16-GMCSF tumors in Sig7/9^+^ mice at different time points following intratumoral administration of Sig7/9*de* (5-15 μg). GMCSF, granulocyte-macrophage colony-stimulating factor. **d**, Assessment of Siglec-7/-9 depletion in tumor-infiltrating myeloid cells (including TAMs, PMN-MDSCs, M-MDSCs and DCs) following administration of 10 μg Sig7/9*de* to B16-GMCSF tumor-bearing Sig7/9^+^ mice for 2 days. **e**,**f**, Assessment of *in vivo* Siglec-7/-9 depletion efficiency in tumor and tumor dLN-infiltrating T cells following administration of 10 µg Sig7/9*de* or PBS to B16-GMCSF tumor-bearing Sig7/9^+^ mice for 2 days. Data are mean ± s.d. Two-tailed unpaired Student’s *t*-test (**a**,**c**,**e**,**f**).

## Siglec-7/9 degradation suppresses tumor growth in syngeneic mouse tumor models

Subsequently, we evaluated the therapeutic potential of Siglec-7/9 degradation in several syngeneic mouse tumor models established by implanting Siglec-7/9L^+^ tumor cell lines (Extended Data Fig. 14a) in Sig7/9^+^ mice. In a B16-GMCSF melanoma model resistant to anti-PD-1 treatment, intratumoral administration of Sig7/9*de* resulted in prolonged survival and notably suppressed tumor growth compared to PBS and non-degrader ((**6**^bio^)_4_-SA-M6P_4_) control. The therapeutical efficacy achieved with Sig7/9*de* is similar to that observed in the SigE^KO^ control group. (Fig. 6a and Extended Data Fig. 14b).

**Fig. 6.**
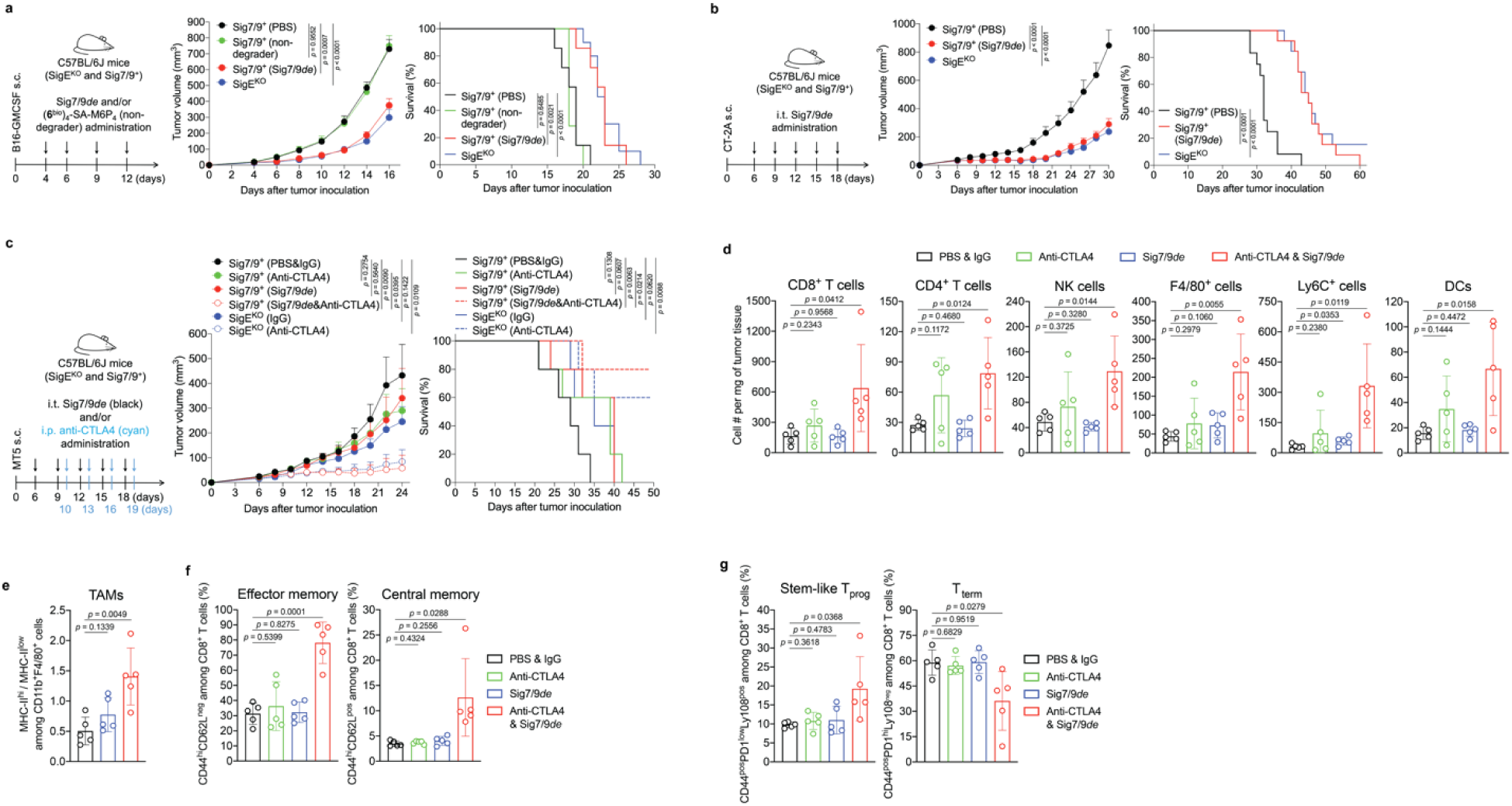
| Siglec-7/9 degradation suppresses tumor growth in syngeneic mouse models. **a**, B16-GMCSF tumor growth and recipient mouse survival in SigE^KO^ (*n*= 11 mice) and Sig7/9^+^ (*n*= 7 mice per group) mice that were intratumorally administrated with PBS, (**6**^bio^)_4_-SA-M6P_4_ and Sig7/9*de*. **b**, CT-2A tumor growth and mouse survival in SigE^KO^ (*n*= 13 mice) and Sig7/9^+^ mice that were intratumorally administrated with PBS (*n*= 12 mice) and Sig7/9*de* (*n*= 13 mice). **c**, MT5 tumor growth and mouse survival in SigE^KO^ (*n*= 5 mice per group) and Sig7/9^+^ (*n*= 5 mice per group) mice that were intratumorally administrated with PBS and Sig7/9*de* from day 6 every 3 days for five doses in total, and/or intraperitoneally administration with anti-CTLA4 from day 10 every 3 days for four doses in total. Average sizes of primary tumors ± SEM are presented in cubic millimeters (mm^3^) (**a**,**b**,**c**). Statistical analysis was performed using one-way ANOVA with Dunnett’s multiple comparisons test (**a**,**b**,**c**). **d**, Flow cytometry analysis of numbers of tumor-infiltrating immune cells in the MT5 tumor model in each treatment condition at day 18 (*n*= 5 mice per group, as described in the methods of **c**). **e**, Analysis of MHC-II^hi^/MHC-II^low^ ratios among tumor-infiltrating TAMs. **f**,**g**, Indicated proportions of effector/central memory (**f**) and progenitor stem-like/terminally differentiated CD8^+^ T cell populations (**g**) among TIL CD8^+^ T cells in the MT5 tumor model in each treatment condition at day 18 (*n*= 5 mice per group, as described in the methods of **c**). Data are mean ± s.d. Two-tailed unpaired Student’s *t*-test (**d**,**e**,**f**,**g**).

Next, the therapeutic efficacy of Sig7/9*de* was further evaluated for treating aggressive glioblastomas (GBM, grade IV glioma), which are often resistant to most checkpoint blockade therapies^2, 34^. A stratification analysis of glioma patients revealed a correlation between the expression of *SEGLEC7* or *SEGLEC9* and reduced overall survival in low-grade gliomas (LGGs) and relapse-free survival in GBMs (Extended Data Fig. 14c,d). Glioma cells express high levels of Siglec-7/9 ganglioside ligands, such as disialoganglioside GD2, GD3 and trisialoganglioside GT1b^35^, which have been used as tumor-associated markers in brain tumors and potential targets for brain cancer therapies^36, 37^. In a subcutaneous CT-2A astrocytoma tumor model with high ganglioside expression^38^, which closely resemble human high-grade gliomas^39^, Sig7/9*de* administration at a 3-day interval effectively inhibited tumor growth and prolonged survival in Sig7/9^+^ mice, in a manner similar to that observed in SigE^KO^ mice (Extended Data Fig. 14e,f and Fig. 6b), suggesting that maximum therapeutic efficacy was achieved. Interestingly, the improved tumor control seemed to be unrelated to macrophage phagocytosis of tumor cells because we did not notice apparent differences in tumor phagocytosis between the Sig7/9*de* treated, untreated, and SigE^KO^ groups (Extended Data Fig. 14g). By contrast, when CD8^+^ T cells were depleted just prior to the start of Sig7/9*de* treatment, the improved tumor control was completely abolished (Extended Data Fig. 14h), suggesting the primary involvement of CD8^+^ T cells in the Sig7/9*de* conferred tumor control.

## Siglec-7/9 degradation synergizes with anti-CTLA4 treatment in a pancreatic cancer model by augmenting TIL stemness and memory repertoire

Pancreatic cancer has a high mortality rate and late diagnosis, which leads to widespread metastasis^40^. PDAC, the most common histologic type of pancreatic cancer, is refractory to anti-CTLA-4 and anti-PD-L1 combination therapy^41, 42, 43^. In the TME of PDAC, TAMs and MDSCs are predominant tumor-promoting players^3^ that block T cell tumor-infiltration and anti-tumor activity^44^, and both are known to be high expressers of Siglec-7/9^9^.

We investigated Siglec-7/9 degradation alone or in combination with anti-CTLA-4 to reprogram the immunosuppressive TME of MT5 pancreatic tumors with mutations recapitulating those found in PDAC patients^45^. Intratumoral administration of Sig7/9*de* led to efficient Siglec-7/9 degradation in tumor-infiltrating immune cells (Extended Data Fig. 15a), but showed only weak tumor control comparable to that induced by anti-CTLA4 alone (i.p.) (Fig. 6c). Interestingly, the co-administration of Sig7/9*de* and anti-CTLA4 substantially suppressed tumor growth and resulted in prolonged overall survival, with 2/5 Sig7/9^+^ mice in the combination treatment group becoming tumor free (Fig. 6c and Extended Data Fig. 15b). These results were similar to those seen with anti-CTLA4 treatment in the SigE^KO^ group (Fig. 6c). Upon rechallenge, these mice quickly rejected the newly introduced tumor cells (Extended Data Fig. 15c), suggesting the formation of immune memory. Furthermore, adoptive transfer of CD8^+^ T cells isolated from tumor-free Sig7/9^+^ mice to naïve Sig7/9^+^ mice demonstrated an excellent ability to control the growth of newly inoculated MT5 tumor cells (Extended Data Fig. 15d,e).

To further explore how Siglec-7/9 degradation synergizes with CTLA-4 blockade to suppress PDAC progression, we conducted a TIL composition and phenotype analysis of MT5 tumors isolated from Sig7/9^+^ mice. Although no apparent changes in the composition of CD4^+^, NK and DCs in tumor dLNs were observed, the single agent and the combination treatment pronouncedly increased the total number of dLN CD8^+^ cells (Extended Data Fig. 15h). Moreover, the combination therapy, but not the monotherapy, also significantly increased the frequency and number of TILs including lymphocytes (CD8^+^ and CD4^+^), NK and myeloid cells (F4/80^+^, Ly6C^+^, and DCs), compared to the PBS control group (Fig. 6d and Extended Data Fig. 15f,g). Although we did not observe altered tumor cell phagocytosis by macrophages (Extended Data Fig. 15i), the combination treatment significantly increased the accumulation of MHC-II^high^ macrophages that are known to possess tumor-suppressive properties in the TME^46^, while decreasing their MHC-II^low^ counterparts (Fig. 6e). In addition, the combination treatment also elicited a remarkable expansion of the memory CD8^+^ T cell population, including effector memory T (T_EM_) and central memory T (T_CM_) cells (Fig. 6f). In the combination treatment group, an increase in the frequency of progenitor CD8^+^ T cells with stem cell-like properties (PD-1^low^, Ly108^+^) was observed with a concomitant decrease in the frequency of terminally differentiated T cells that are PD-1^hi^ Ly108^−^(Ly108 serves as the surrogate marker for TCF-1, the transcription factor essential for T cell stemness^47, 48^) (Fig. 6g). Taken together, these observations provide strong evidence that the combination treatment converts the immunosuppressive TME with poor T-cell infiltration to a relatively permissive T-cell enriched TME that is sensitive to immune checkpoint blockade.

## Discussion

In healthy humans, Siglec-7 and -9 are predominantly expressed by myeloid cells with very low expression on normal T cells^49, 50, 51^. However, upregulation of Siglec-9 has been observed on tumor-infiltrating T cells in patients with colorectal, ovarian cancer, and melanoma^49^. Likewise, Siglec-7 and -9 are upregulated on tumor-infiltrating T cells in non–small cell lung cancer (NSCLC) patients^49^. By analyzing tumor samples from PDAC patients (Fig. 1c,d), we also observed high expression of Siglec-7 and -9 on T cells, despite their disproportionately low mRNA levels (Fig. 1a,b). The origin of these Siglecs on T cells was puzzling. We discovered by serendipity that in the complex TME, T cells readily acquire these inhibitory Siglec molecules from the neighboring myeloid cells, resulting in suppressed T cell activation and effector function (Extended Data Fig. 16). These findings suggest that scRNA-seq, not alone but in combination with complementary spatial analysis techniques, may provide a more comprehensive and unbiased understanding of cell-cell interactions and outcomes. These findings also highlight the importance of considering not only intrinsic, but also extrinsic checkpoints acquired by T cells from specific TMEs when selecting checkpoint blockade therapy to reinvigorate T-cell immunity.

To inhibit the immunosuppressive Siglec-sialoglycan interactions, two complementary approaches have been pursued: direct inhibition of Siglecs using functional blocking antibodies^14,52^ and targeted desialylation^7, 31^. For Siglec-7 and -9, while blocking their interaction with sialoglycans was beneficial^14^, both commercial and custom-made anti-Siglec-9 antibodies also trigger inhibitory signaling due to their agonistic nature, as exemplified in ref. (49) and Extended Data Fig. 4d. Targeted desialylation has been achieved by selectively removing sialoglycans using antibody-sialidase conjugates^7, 31^ and bispecific T cell engager-sialidase fusion proteins^29^. However, similar to other antibody or CAR-T cell-based immunotherapies, only a subset of tumor cells expressing the target antigen can be desialylated due to the antigenic heterogeneity of solid tumors.

Using SuFEx click chemistry we have developed a specific degrader to induce targeted lysosomal degradation of both Siglec-7 and -9. Compared to antibody/nanobody-based extracellular targeted protein degraders^53, 54^ that induce only partial degradation of engaged membrane proteins in cultured cells, Sig7/9*de* exhibits better efficacy, as nearly quantitative degradation of Siglec-7/9 was observed in both cell lines and primary cells. Although complete degradation was not observed *in vivo* (% of degradation = 50-70%), efficient tumor control was achieved in all tumor models with the therapeutic effect comparable to that observed in SigE^KO^ mice. Thus, Sig7/9*de* represents the first agent capable of inducing simultaneous degradation of two membrane receptors to achieve maximum therapeutic benefit.

Siglec-7/9 receptors are abundantly expressed on TAMs, which play diverse roles within the TME, including regulation of inflammation and modulation of adaptive immune responses. Rather than TAM depletion, which may disrupt this delicate balance, potentially leading to dysregulated immune responses and adverse effects^55^, Siglec-7/9 degradation leads to TAM and T cell reprogramming, resulting in effective tumor control. In contrast to observations in SigE^KO^ mice, in which Siglec-E deletion was found to facilitate tumor cell phagocytosis by microglia and monocyte-derived cells^56^, Siglec-7/9 degradation had a negligible effect on macrophage-mediated tumor cell phagocytosis, but enhanced antigen-presentation capability of TAMs when combined with CTLA-4 blockade.

Several human Siglecs have been identified as immunosuppressors in the TME, including Siglec-7, -9, -10 and -15^6, 14, 52, 56, 57, 58, 59^, but their relative contributions to the cancer-immunity cycle in different tumor types remain to be explored. As discovered by Bertozzi and coworkers, the therapeutic effects of targeted desialylation *in vivo* are largely dependent on functional Siglec-E expression — no discernible anti-tumor benefit was observed for their αHER2 antibody– sialidase conjugate in SigE^KO^ mice^7, 31^. This observation underscores the need to develop therapeutics targeting multiple Siglecs that play dominant roles in the microenvironment of certain tumors, such as the Siglec-7/9 degrader described in this study, as well as Siglec-targeted therapeutics with broad neutralizing activity.

## Methods

### Cell culture

All cells were grown at 37 °C in a humidified incubator under 5% CO_2_. U937 cells and their variants, including Siglec-7/9 KO, Siglec-7 WT, Siglec-7 R124A, Siglec-9 WT and Siglec-9 R120A, were grown in RPMI-1640 supplemented with GlutaMAX, 10% fetal bovine serum (FBS) and 1% penicillin/streptomycin (P/S), and cell density was maintained between 0.1 x 10^6^ and 1 x 10^6^ viable cells/mL. Jurkat cells and their variants, including EV (empty vector), Siglec-7 WT and Siglec-9 WT, were grown in RPMI-1640 supplemented with GlutaMAX, 10% FBS, 1% P/S, 1 mM sodium pyruvate, 10 mM HEPES, and 1% non-essential amino acids (NEAA). Primary human and mouse T cell activation and expansion was performed in RPMI-1640 supplemented with GlutaMAX, 10% FBS, 1% P/S, 1 mM sodium pyruvate, 10 mM HEPES, 1% NEAA and 50 μM β-mercaptoethanol. CHO (Chinese hamster ovary) cells expressing Siglecs (WT and R-mutant) and Siglec-Fcs were grown in DMEM/F-12 supplemented with 10% FBS, 1% P/S, and 15 mM HEPES. HEK293T cells (for lentiviral vector production) and Plat-E cells (for retroviral vector production) were grown in DMEM supplemented with 10% FBS and 1% P/S. Ramos, T47D, OVCAR3, SF295, CCRF-CEM and MT5 cells were grown in RPMI-1640 supplemented with GlutaMAX, 10% FBS and 1% P/S and 1% NEAA. MDA-MB-231, HT29, SUIT2, B16, B16-GP33, B16-GMCSF, CT-2A cells were grown in DMEM supplemented with 10% FBS and 1% P/S and 1% NEAA.

B16-GMCSF cell line was a gift from Dr. Jonathan Kagan (Harvard Medical School). CT-2A cell line was purchased from Dr. Thomas Seyfried (Boston College). SUIT2 and MT5 cell lines were gifts from Dr. David Tuveson (Cold Spring Harbor Laboratory).

### CRISPR/Cas9 genetic knockout of Siglec-7/-9 in U937 cells

Double knockout (KO) of Siglec-7 and -9 in U937 cells was performed using the CRISPR/Cas9 as described^60^. This approach involved sequential KO of each Siglec individually, resulting in a Siglec-7/-9 double KO (Sig7/9-KO) phenotype in the U937 cell line.

### Siglec-7/-9 WT and R-mutant transduction

Lentiviruses encompassing Siglec-7 WT and Siglec-9 WT, as well as their R-mutants were produced by polyethylenimine (PEI)-based transfection of HEK293T cells using Siglec-expressing RP172^60^, pMD2G, pRSV-Rev and pMDLg/p. For lentiviral transduction, 5×10^5^ Sig7/9-KO U937 cells or Jurkat cells were seeded in 800 μL complete medium supplemented with 8 μg/mL polybrene (Sigma-Aldrich) and lentivirus. After 1 day, the medium was topped up to 2 mL. The cells were expanded for additional 3 days and Siglec^+^ population was sorted by staining cells with anti-Siglec-7 APC (BioLegend, clone 6-434) or anti-Siglec-9 APC (BioLegend, clone K8).

### Retroviral transduction of Siglec-7/-9 in P14 T cells

Retroviruses were generated by transfecting Plat-E cells with pMIGR1 vectors expressing EV, Siglec-7 or Siglec-9, using PEI. Supplementary Table S1 and S2 provide primers for cloning of Siglec-7 and -9 into pMIGR1. For the production of retroviruses containing both Siglec-7 and -9, vectors were added in a 1:1 ratio. After transfection for 6 hours the cell medium was refreshed, and viral supernatants were harvested 48 hours later. The supernatants were filtered, concentrated using Retro-X™ Concentrator (Takara Bio) and stored in aliquots at – 80 °C. For retroviral transduction of P14 T cells, P14 splenocytes at 1×10^6^/mL density were first activated with 10 nM GP33 peptide for two days. The activated cells, identified as P14 CD8^+^ T cells with > 98% purity, were then spinfected (2000*g*, 2 hours, 32 °C) with virus containing single Siglec-7 or -9, or a combination of both Siglecs. After overnight culture with 100 U/mL IL2, the Siglec^+^ population was sorted by staining the cells with anti-Siglec-7 APC (BioLegend, clone 6-434) or anti-Siglec-9 APC (BioLegend, clone K8). The sorted cells were expanded with 100 U/mL IL2 for 4-7 days.

### U937 cell differentiation

U937 cells and their variants in complete medium at 5×10^5^/mL density were treated with 10 nM phorbol 12-myristate 13-acetate (PMA) for 2 days followed by washing twice with PBS. The adherent cells were rested in fresh complete medium for another 2 days.

### hMDM generation and stimulation

Anonymous healthy donor blood samples were obtained from Scripps Research’s Normal Blood Donor Services (NBDS). PBMCs were isolated thought density gradients using Ficoll. Monocytes were then enriched from PBMCs using Human CD14 Positive Selection Kit II (Stemcell Cat # 17858). The enriched monocytes were differentiated into macrophages in IMDM (Gibco, Cat# 12440053) supplemented with 10% FBS and exogenous cytokines by 7–9 days of culture. To generate M0, monocytes were cultured in the presence of 50 ng/mL human M-CSF (BioLegend) for 7-9 days; to generate M2, monocytes were cultured in the presence of 50 ng/ml M-CSF for 4– 5 days, followed by incubation with 50 ng/mL M-CSF, 50 ng/mL IL10 (Genscript) and 50 ng/mL TGFbeta (Genscript) in fresh medium for another 2-4 days.

### Mouse BMDM and BMDC generation

Bone marrow cells were flushed from tibias and femurs using a syringe into IMDM medium supplemented with 10% FBS and 1% P/S. Cells were collected, and red blood cells (RBCs) were lysed in ACK lysis buffer. The remaining cells were then plated at 1×10^6^/mL density in IMDM medium containing 20 ng/mL mouse M-CSF (BioLegend) and cultured for 7-9 days without medium change to generate BMDMs.

For BMDC production, bone marrow cells were plated at 3×10^5^/mL density in IMDM medium containing 20 ng/mL mouse GM-CSF (BioLegend) and cultured for 7-9 days.

### Siglec-Fc preparation

Siglec-Fc expressing CHO cells^61^ were seeded in a T75 flask in 30 mL DMEM/F-12 supplemented with 1.5% FBS, 1% P/S, and 15 mM HEPES. The supernatant was collected after 1-week culture upon full confluency, centrifuged at 1000*g* for 10 min and filtered to remove debris. Protein A agarose beads (Thermo Fisher Scientific, Cat#: 15918014) was washed with PBS and incubated with the cleared supernatant at r.t. for 2 hours. The resulting mixture of supernatant plus beads was applied to a disposable column, washed with 4 mL sodium phosphate buffer (20 mM, pH 7.4), eluted with glycine buffer (120 mM, pH 2-3), and neutralized with Tris buffer (pH 9). The neutralized eluates containing protein fractions were pooled, concentrated and buffer-exchanged to sterile PBS. The final products were stored in aliquots at – 80 °C.

### Cell-surface screening assay

Jurkat cells lacking the expression of Siglec receptors were used to install synthetic Neu5Ac mimetics on the cell surface in a natural and multivalent context for high-affinity Siglec ligand discovery. Briefly, Jurkat cells were first fed with 50 μM Ac_4_ManPoc for 3 days in complete medium for preparing alkynalylated cells through metabolic glycoengineering. Then, the alkynalylated cells were seeded to 96-well microplates (2×10^5^ cells per well) in PBS buffer containing 1% FBS, pre-mixed CuSO_4_/BTTPS (75 μM/450 μM, 1:6) and synthetic Neu5Ac-LacNAc-azide ligands (200 μM). Treatment with sodium ascorbate (2.5 mM) at room temperature (r.t.) for 15 min was used to initiate the BTTPS-accelerated CuAAc, followed by quenching with 1 mM bathocuproine disulfonate (BCS), during which cell viability was maintained after washing with PBS. Finally, the resulting Neu5Ac-LacNAc mimetics installed on the cell surface were probed with pre-mixed recombinant Siglec-Fc chimera and anti-Fc FITC or APC for FACS analysis of the indicated cells for comparison of mean fluorescence intensity (MFI) signals.

### Siglec-7/9 ligand tetramer preparation

(SigL^bio^)_4_-SA tetramers were prepared by combining biotinylated Siglec ligands with SA (with or without modification with HRP, fluorophore, or M6P) in a molar ratio of 4:1 at 4 °C overnight. The (SigL^bio^)_4_-SA-M6P_4_ tetramer was utilized as either a degrader ((**8**^bio^)_4_-SA-M6P_4_) or a non-degrader ((**6**^bio^)_4_-SA-M6P_4_).

### ELISA-like assay for measuring Siglec ligand cross reactivity

96-well high-binding microplates (Corning 9018) were coated with 5 μg/mL protein A in coating buffer at 4 °C overnight. After washing and blocking, the plates were incubated with 5 μg/mL Siglec-Fc chimeras or human IgG at r.t. for 2 hours, followed by incubation with (SigL^bio^)_4_-SA-HRP tetramers at r.t. for 1 hour. The colorimetric HRP substrate (TMB) was added into each well at r.t. for 15 min, and the reaction was stopped by addition of 1 M H_3_PO_4_. The absorbances of each well were analyzed at 450 nm using a plate reader.

### Biolayer Interferometry (BLI) assay

The BLI assay was performed on an Octet Red96 (ForteBio) instrument. Recombinant Siglec-7-Fc and Siglec-9-Fc were loaded onto Octet AHC2 biosensors (Satorius, part #18-5142) at concentrations of 10 μg/mL and 5 μg/mL, respectively, in kinetics buffer (0.05% Tween-20 in PBS) for 2 min. Association of (**8**^bio^)_4_-SA and (**9**^bio^)_4_-SA tetramers was conducted by immersing the biosensors in the kinetic buffer at concentrations ranging from 25 to 1000 nM for 3 min. Association of anti-Siglec-7 (BioLegend, clone S7.7) and anti-Siglec-9 (BioLegend, clone K8) was conducted by immersing the biosensors in the kinetic buffer at concentrations ranging from 5 to 200 nM for 3 min. Dissociation was recorded in the kinetics buffer for 5 min. The recorded signals were corrected by subtracting the reference background and the *K*_D_, *k*_on_ and *k*_off_ values were calculated using a global fit model with the Octet Data Analysis software.

### Siglec staining with Siglec ligand tetramers and antibodies

Cells expressing Siglec KO, WT or R-mutant (∼ 1×10^5^) were suspended in FACS buffer (PBS containing 1 mM EDTA and 1% FBS) and incubated with 1 μg/mL (SigL^bio^)_4_-SA-AF647 or 1 μg/mL anti-Siglec antibody at 4 °C for 30 min, followed by washes and FACS analysis.

### Siglec ligand staining

#### Detection of Siglec ligands on mammalian cancer cells

Recombinant Siglec-Fc (5 μg/mL) and APC anti-IgG Fc (2.5 μg/mL) were pre-mixed in FACS buffer (PBS containing 0.5 mM EDTA and 1% FBS) for 30 min on ice. Adherent cancer cells were trypsinized and collected using TrypLE Express. ∼1×10^5^ trypsinized cells or non-adherent cells were suspended in 50 μL of the above FACS buffer containing the pre-mixed Siglec-Fc/anti-IgG APC and incubated on ice for 40 min. Cells were washed twice with FACS buffer, followed by flow cytometry analysis.

#### Detection of Siglec ligands on donor PBMC and mouse splenic T cells

Donor PBMCs and mouse splenocytes were first blocked with human Fc blocker (BioLegend, TruStain FcX, Cat# 422302) and mouse Fc blocker (BioLegend, TruStain FcX PLUS, Cat# 156603) respectively, in FACS buffer (PBS containing 0.5 mM EDTA and 1% FBS). Cells were then incubated with the pre-mixed Siglec-Fc/anti-IgG APC in FACS buffer on ice for 30 min. After that, fluorescent antibodies including anti-CD3, anti-CD8, anti-CD4 and anti-CD11b were added at 1:200 dilution and incubated on ice for further 30 min. Finally, cells were washed and analyzed by gating on T cells using flow cytometry.

### Immune cell staining with anti-CI-M6PR

Immune cells, including U937, Jurkat, hMDMs, P14 T cells and BMDMs, were suspended in 50 μL FACS buffer, blocked with Fc blocker if needed, and incubated with primary anti-CI-M6PR (Abcam, clone 2G11) on ice for 30 min. Cells were washed twice with FACS buffer and incubated with anti-mouse IgG AF488 or APC in 50 μL FACS buffer, followed by CI-M6PR expression analysis using flow cytometry.

For intracellular CI-M6PR staining, cells were first fixed, permeabilized and Fc blocked, then stained with anti-CI-M6PR and fluorescent anti-mouse IgG, followed by flow cytometry analysis.

### Cancer cell CD47 staining

Approximately 1×10^5^ trypsinized adherent or non-adherent cancer cells were suspended in 50 μL FACS buffer containing the primary anti-CD47 (BioXCell, clone B6H12), and incubated on ice for 30 min. Cells were washed twice with FACS buffer and incubated with anti-mouse IgG APC in 50 μL FACS buffer, followed by CD47 expression analysis using flow cytometry.

### Siglec recycling experiments

#### Detection of Siglec internalization by flow cytometry

Siglec-7 and Siglec-9 expressing U937-derived macrophages were trypsinized and collected using TrypLE Express (Gibco, Cat# 12604013). Siglec-7^+^ and Siglec-9^+^ macrophages in complete medium were treated with 100 nM (**8**^bio^)_4_-SA-AF488 in an ultra-low attachment plate at 37 °C. Ice-cold FACS buffer was added at various time points (5, 15, 30, 60 and 120 min), then cells were washed twice and stained with anti-Siglec-7 APC (BioLegend, clone 6-434) or anti-Siglec-9 APC (BioLegend, clone K8) on ice for 30 min, followed by flow cytometry analysis of AF488 uptake and cell-surface Siglec levels. Washing and staining procedures were always performed on ice.

#### Detection of Siglec internalization by microscopy

2.5×10^5^ trypsinized Siglec-7/9 KO and WT macrophages in 0.5 mL complete medium were seeded on a sterile #1.5 coverslide (10 mm) in a 24-well plate overnight. Cell medium was replaced with fresh medium treated with 100 nM SA-AF488 or 100 nM (**8**^bio^)_4_-SA-AF488 tetramer. After incubation at 37 °C for 30 min, cells were fixed with 4% paraformaldehyde (PFA) in PBS at r.t. for 20 min, permeabilized with 0.25% Triton X-100 in PBS at r.t. for 13 min. Cells were then washed with PBST buffer (0.1% Tween-20 in PBS) and blocked with 30 μg/mL hIgG (polyclonal human IgG; R&D Cat# 1-001-A) in 4% FBS in TBST at r.t. for 60 min. After that, cells were incubated with goat anti-Siglec-7 or -9 (R&D AF1138 and AF1139, respectively) and Rb anti-Rab5 (Cell Signaling, clone C8B1, Cat# 3547) diluted in 4% FBS in PBST containing 5 μg/mL hIgG at 4 °C overnight. Cells were washed with PBST buffer and incubated with secondary AF555 Dnk anti-goat and AF647 Dnk anti-Rb (Abcam) diluted in 4% FBS in PBST containing 5 μg/mL hIgG at r.t. for 1 hour (protected from light). Cells were washed with PBS and incubated with 1.5 µg/mL DAPI in PBS at r.t. for 10 min, followed by mounting coverslides onto microscope slides in one drop of anti-fade fluorescence mounting medium (Invitrogen, Cat# P36961). The imaging was measured at x60 oil immersion objective using Zeiss LSM 780 confocal laser scanning microscope at Scripps Core Microscopy Facility, and image analysis was performed using ImageJ.

#### Detection of Siglec recovery by microscopy

2.5×10^5^ trypsinized Siglec-7^+^ and Siglec-9^+^ macrophages in 0.5 mL complete medium were seeded on a sterile #1.5 coverslide (10 mm) in a 24-well plate overnight. Cell medium was replaced with fresh medium treated with PBS or 100 nM (**8**^bio^)_4_-SA-AF488 tetramer. After incubation at 37 °C for 30 min, cells were washed three times with PBS, and covered with 0.5 mL fresh complete medium. After different time points (0, 3, 6, 12 and 24 hours), cells were fixed with 4% PFA in PBS at r.t. for 20 min, followed by permeabilization with 0.25% Triton X-100 in PBS at r.t. for 13 min. Cells were washed, blocked, stained with primary antibodies followed by secondary antibodies, and imaging was measured and analyzed as described above.

#### Detection of Siglec recovery by flow cytometry

Trypsinized Siglec-7^+^ and Siglec-9^+^ macrophages in complete medium were treated with PBS or 100 nM (**8**^bio^)_4_-SA-AF488 tetramer, in an ultra-low attachment plate at 37 °C for 30 min. Ice-cold PBS was added, and cells were washed three times with PBS, resuspended in complete medium, and seeded in the ultra-low attachment plates. After different time points (0, 3, 6, 12 and 24 hours), cells were harvested, washed and stained with anti-Siglec antibodies (R&D, AF1138 and AF1139) for analysis of cell-surface Siglec recovery. Washing and staining were always performed on ice.

### Siglec degradation

#### Western blot experiment

Differentiated U937-derived macrophages (2.5×10^5^ cells) in 0.5 mL RPMI complete medium, or hMDMs (8×10^4^ cells) in 0.5 mL IMDM complete medium, or BMDMs (1.5×10^5^ cells) in 0.5 mL IMDM complete medium were seeded in 24-well plates for 1 day. The cell medium was replaced with fresh 0.5 mL medium treated with (**8**^bio^)_4_-SA-M6P_4_ (Sig7/9de), (**8**^bio^)_4_-SA, SA-M6P_4_ or PBS ctr for the indicated periods. For degradation inhibitor treatment, 50 nM bafilomycin A1 (BafA1) or 10 µM MG132 was added to the macrophages for 30 min prior to Siglec degrader treatment. After that, cells were washed three times with cold PBS, and lysed with 80 μL RIPA buffer (Alfa Aesar, J63306) supplemented with protease inhibitor cocktail (Roche), phosphatase inhibitor cocktail (Cell Signaling Technologies) and 5 μg/mL DNase I on ice for 50 min. Cell lysates were transferred to microcentrifuge tubes and centrifuged at 15,000g for 10 min at 4 °C. The supernatant was collected, and the protein concentration was quantified by bicinchoninic acid (BCA) assay (Pierce). Equal amounts of lysates were resolved by SDS-PAGE (11% acrylamide), then transferred to a nitrocellulose membrane by semi-dry electrophoretic transfer. The membrane was blocked with 5% BSA in Tris-buffered saline with 0.05% Tween-20 (TBST) buffer at r.t. for 1 hour with shaking. Membranes were incubated with primary antibodies (0.1 µg/mL dilution for both anti-Siglec-7 (R&D, AF1138) and anti-Siglec-9 (R&D, AF1139); 1:1000 dilution for anti-β-actin (BioLegend) with fresh 5% BSA in TBST buffer) at 4 °C overnight with shaking. Membrane was washed four times with TBST buffer and incubated with secondary antibody (1:5,000 dilution for both anti-mouse IgG HRP and anti-goat IgG HRP with fresh 5% BSA in TBST buffer) at r.t. for 1 hour with shaking. After washing with TBST buffer for four times, the protein signals in the membrane were visualized and recorded using SuperSignal West Pico PLUS Chemiluminescent Substrate by Bio-Rad chemiluminescence system. Quantification of relative band intensities was processed using ImageJ.

#### Cell surface degradation experiment

For adherent cells, macrophages in 0.5 mL complete medium in 6- or 24-well plates were treated with Sig7/9de for the indicated periods, then trypsinized and collected using TrypLE Express. The trypsinized macrophages were stained with anti-Siglec-7 APC (BioLegend, clone 6-434) or anti-Siglec-9 APC (BioLegend, clone K8) for flow cytometry analysis of cell-surface Siglec levels. Staining and washing steps were always performed on ice using cold FACS buffer. Alternatively, macrophages were first trypsinized and collected using TrypLE Express. Trypsinized macrophages in 0.5 mL complete medium were then seeded in 24-well flat-bottom ultra-low attachment plates (Corning 3473) for degrader treatment, followed by anti-Siglec staining for flow cytometry analysis in the same procedure. For non-adherent cells such as Siglec^+^ Jurkat and P14 T cells, 5×10^4^ cells were suspended in 100 μL RPMI complete medium, seeded in 96-well plates, treated with degrader and stained with anti-Siglec antibodies for flow cytometry analysis.

#### Microscope-measured degradation experiment

2.5×10^5^ trypsinized Siglec-7/9 KO and WT macrophages in 0.5 mL complete medium were seeded on a sterile #1.5 coverslide (10 mm) in a 24-well plate overnight. Cell medium was replaced with fresh medium treated with PBS or 30 nM Sig7/9de. After 24 hours, cells were fixed with 4% PFA in PBS at r.t. for 20 min, permeabilized with 0.25% Triton X-100 in PBS at r.t. for 13 min. Cells were then washed, blocked, stained with goat anti-Siglec-7 or -9 (R&D AF1138 and AF1139, respectively), followed by staining with AF555 Dnk anti-goat (Abcam) and DAPI, and imaging was measured and analyzed as described above.

### Cancer phagocytosis

#### Flow cytometry-based assay

IL10/TGFβ-polarized M2 phenotypic hMDMs with or without Siglec-7/9 pre-degradation were trypsinized and collected using TrypLE Express. Siglec-7/9 were pre-degraded in the indicated hMDMs by treatment with 30 nM Sig7/9de for 24 hours in the incubator. Cancer cells were also trypsinized and collected using TrypLE Express, followed by labeling with 1 μM calcein-AM (BioLegend) in PBS at 37 °C for 15 min. hMDMs and ‘green’-labeled cancer cells were cocultured at a 1:2 ratio in the presence or absence of 10 μg/mL antibodies or IgG control in serum-free IMDM medium in the 96-well round-bottom ultra-low attachment plates (Corning 7007) for 3 hours in the incubator. In the coculture setup, cancer cells were pre-opsonized with anti-CD47 (BioXCell, clone B6H12) at 37 °C for 30 min before coculture with hMDMs. After coculture, phagocytosis was halted by the addition of ice-cold FACS buffer. Cells were centrifuged at 500g for 4 min, Fc blocked and stained with anti-CD11b APC (BioLegend, clone ICRF44) in FACS buffer on ice for 30 min. Cells were then stained with 1 μg/mL DAPI in PBS for 10 min followed by flow cytometry analysis. Phagocytosis index was quantified as the number of phagocytosing macrophages (CD11b^+^ calcium-AM^+^) / total number of the live CD11b^+^ population per 100 macrophages after the removal of debris and doublets. Each phagocytosis assay was performed with a minimum of three technical replicates.

#### Microscopy-based assay

IL10/TGFβ-polarized M2 hMDMs (with or without Siglec-7/9 pre-degradation) and HT29 cells were trypsinized and collected using TrypLE Express. HT29 cells were washed twice with PBS and incubated with 5 μM pHrodo Red-succinimidyl ester (Invitrogen, Cat# P36600) with rotation at r.t. for 30 min in the dark. After pHrodo labeling, HT29 cells were washed with complete medium and opsonized with anti-CD47 or IgG control for 30 min. In the meantime, hMDM were labeled with 4 μM CFSE (BioLegend, Cat# 423801) in PBS at r.t. for 5 min, and washed with complete medium. 3×10^4^ hMDMs and 9×10^4^ HT29 cells were mixed in 60 μL serum-free IMDM medium and seeded in a 96-well flat-bottom tissue-treated plate. After 3-hour coculture in the incubator, cells were fixed with 4% PFA in PBS at r.t. for 20 min and gently washed twice with PBS. The phagocytic red signal and macrophage green signal were automatically acquired at x20 objective at 300 ms (green) and 600 ms (red) exposures per field using the Nikon Ti2-E inverted motorized microscope at Scripps Core Microscopy Facility. Each phagocytosis reaction was carried out in four replicates, and reported values are averaged over four distinct fields per well. The phagocytosis index was calculated as described above.

### Siglec trogocytosis

#### Confocal microscopic analysis of immunological synapse

hMDM-PBMC T cell immunological synapse was captured for Siglec-7/9 trogocytosis study. PBMC T cells were isolated from healthy donor using a MojoSort human CD3 T cell isolation kit (BioLegend), and FACS-sorted to yield the Siglec-7/9^−^ population for subsequent coculture with macrophages. Autologous hMDMs were trypsinized and collected using TrypLE Express, followed by pulsing with 1 μg/mL SEB at 37 °C for 2 hours in a 24-well ultra-low attachment plate. Then, 2×10^5^ SEB-pulsed hMDMs and 5×10^5^ Siglec-7/9^−^ T cells in each 80 μL IMDM complete medium, were mixed and seeded in a 96-well round-bottom ultra-low attachment plate, followed by centrifugation at 150g for 1 min to initiate the cell-cell interactions. After coculture at 37 °C for 30 min, cells were gently fixed with 4% PFA in PBS at r.t. for 20 min and permeabilized with 0.25% Triton X-100 in PBS at r.t. for 11 min. After blocking, cells were stained with goat anti-Siglec-7 or -9 (R&D AF1138 and AF1139, respectively) and Rb anti-CD3ε (Cell Signaling, D7A6E, Cat# 85061T), followed by staining with secondary antibodies (AF555 Dnk anti-goat and AF647 Dnk anti-Rb). Finally, cells were loaded into an 8-well chambered coverslides (Lab-Tek II chambered #1.5 German coverglass system) in PBS for confocal imaging using x60 oil immersion objective as described above.

#### Flow cytometry analysis of Siglec trogocytosis

3×10^4^ PBMC T cells and 1.5×10^4^ autologous hMDMs in each 50 μL IMDM complete medium were cocultured as described above. Mouse splenic T cells enriched using the mouse T cell isolation kit (STEMCELL) and SEB-pulsed SigE^KO^ or Sig7/9^+^ BMDMs (or BMDCs) were seeded at 2:1 ratio (T : BMDM or BMDC) and cocultured in the same manner. After coculture at 37 °C for the indicated periods, cells were harvested, washed and blocked with Fc blocker, then stained with anti-CD3 AF700, anti-CD11b FITC, anti-Siglec-7 PE and anti-Siglec-9 APC, followed by DAPI (for human T cell analysis); or stained with anti-CD45.1 FITC, anti-CD45.2 PE/Cy7, anti-Siglec-7 PE and anti-Siglec-9 APC, followed by DAPI (for mouse T cell analysis).

For T cell incubation with hMDM culture medium, the medium was collected from the supernatant of hMDM cell culture followed by centrifugation at 500g for 5 min.

For the transwell experiment (0.4 μm filter, Corning 3381), macrophages were seeded in the transwell insert (upper filter) and T cells were placed in the lower chamber.

#### Western blot analysis of Siglec transfer

1.5×10^6^ B6 splenic T cells and 7.5×10^5^ Sig7/9^+^ BMDMs or SigE^KO^ BMDMs were cocultured in 200 μL IMDM medium in a 96-well round-bottom ultra-low attachment plate for 30 min. After that, T cells were isolated using mouse T cell isolation kit to remove the BMDMs. Enriched T cells and BMDM control cells were washed twice with PBS and lysed with 40 μL RIPA lysis buffer (containing protease and phosphatase inhibitor cocktails) on ice for 1 hour. Cell lysates were cleared by centrifugation at 15,000g for 10 min at 4 °C, Equal amounts of supernatants were subjected to SDS-PAGE (11% acrylamide) and transferred to a nitrocellulose membrane by semi-dry electrophoretic transfer. The membrane was blocked and probed with anti-Siglec-9 (R&D, AF1139), anti-Siglec-7 (R&D, AF1138), and anti-β-actin (BioLegend) as described above.

#### Immunostaining of Siglecs on tumor tissue T cells

Formalin-fixed paraffin-embedded (FFPE) PDAC tissue slides prepared from grade II and III patients were purchased from TissueArray.Com. Paraffin was first removed by immersing the slides in 2 changes of xylene. Tissue samples were then rehydrated by sequential immersion in 100% ethanol, 95% ethanol, 70% ethanol, 50% ethanol and ddH_2_O. Antigen retrieval was performed by boiling slides in citrate buffer (pH 6) for 20 min, followed by immersion in cold PBST buffer. After that, samples were blocked and stained with goat anti-Siglec-7 or -9 (R&D AF1138 and AF1139, respectively), Rb anti-CD3ε (Cell Signaling, D7A6E, Cat# 85061T), and mouse anti-pan-cytokeratin (Novus Biologicals, AE-1/AE-3, Cat# NBP2-29429). Finally, samples were stained with secondary antibodies (AF555 Dnk anti-goat, AF647 Dnk anti-Rb and AF488 Dnk anti-mouse) and DAPI, followed by mounting coverslides onto the slides for imaging using x20 oil immersion objective as described above.

#### In vivo Siglec trogocytosis

Sig7/9^+^ mice were inoculated with B16-GP33/GMCSF tumors as described in **Tumor models and treatments**. On day 5, CD45.1^+/–^ P14 T cells (prepared as described above) in 100 μL serum-free medium were adoptively transferred into the B16-GP33/GMCSF bearing mice (2×10^6^ T cells per mouse). On day 9, tumors, tumor dLNs, spleens and blood samples were harvested for the analysis of Siglec-7/9 appearance on P14 and endogenous CD8^+^ T cells by staining with anti-CD45.1 AF700, anti-CD45.2 FITC, anti-CD8a PE/Cy7, anti-CD11b PerCP/Cy5.5, anti-NK1.1 PB, anti-Siglec-7 PE, and anti-Siglec-9 APC.

For Siglec-7/9^+^ T cell adoptive transfer, CD8^+^ T cells containing ∼ 75% Sig7/9^+^ population were isolated from a Sig7/9^+^ mouse spleen and labeled with CFSE. 4×10^6^ CFSE^+^ CD8^+^ T cells in 100 μL serum-free medium were i.v. injected to each SigE^KO^ mouse. On days 1 and 4, spleens and inguinal LNs were harvested for the analysis of Siglec-7/9 expression on the adoptively transferred CFSE^+^ T cells.

For SigE^KO^ T cell adoptive transfer, CD8^+^ T cells were isolated from a SigE^KO^ mouse spleen and labeled with CFSE. 3×10^6^ CFSE^+^ CD8^+^ T cells in 100 μL serum-free medium were i.v. injected to Sig7/9^+^ mice. After 1 day, spleens were harvested for the analysis of Siglec-7/9 presence on the adoptively transferred CFSE^+^ and endogenous T cells.

### Quantitative PCR

Total RNA was extracted and purified from splenic CD11b^+^ and T cells isolated from SigE^KO^ and Sig7/9^+^ mice, respectively. Polymerase chain reaction (PCR) was carried out in 10 μL in triplicate using the iTaq Universal SYBR Green One-Step kit (Bio-Rad, Cat# 1725150) according to the manufacturer’s instructions, on a QuantStudio^TM^ 6 Pro System (Thermo Fisher Scientific) at Scripps Biophysics and Biochemistry Core. *SIGLEC* expression was analyzed by the 2^-ΔΔCt^ method, wherein the Ct values of *SIGLEC* genes were normalized to the Ct values of mouse *GAPDH* (glyceraldehyde-3-phosphate dehydrogenase). Supplementary Table S3 lists the primers used for PCR.

### Public scRNA-seq dataset analysis of *SIGLEC7/9*

The expression of *SIGLEC7* and *SIGLEC9* in cell types of the tumor microenvironment was analyzed by examining the following publicly available scRNA-seq datasets of human solid tumors: glioma (Study 1: GEO accession GSE192109, Study 2: Broad Single Cell Portal accession SCP2389), breast cancer (EGA accession EGAS00001005115) and colon cancer (GEO accession GSE178341). Cell type assignments provided by the authors were used. The expression was analyzed and plotted using the Broad Institute Single Cell Portal using the following parameters: subsampling of up to 100,000 cells, annotation by Assignment (GSE192109), Annotation (SCP2389), Cell Type (EGAS00001005115) and ClusterTop (GSE178341) with outliers displayed as individual dots on the violin plots. scRNA-seq PDAC data integrated previously (PMID: 37633924) were analyzed using Seurat v5.0.1. The DotPlot function was used for plotting expression levels. Cell type annotations provided by the authors were used (PMID: 37633924).

### T cell response assays

#### Ex vivo restimulation of P14 T cells in tumor and tumor dLNs

Thy1.1^+/–^ P14 T cells were first activated with 10 nM GP33 peptide for two days as described above, and then adoptively transferred into Sig7/9^+^ mice inoculated with B16-GP33/GMCSF tumors (2×10^6^ cells per mouse as described in **Tumor models and treatments**). After 4 days, cell suspensions from whole tumor and tumor dLN were prepared respectively and plated in 24-well flat-bottom plates, followed by treatment with 100 pM GP33 peptide. After incubation at 37 °C for 1 hour, 1x brefeldin A (BioLegend) was added to the cell suspensions. After further incubation at 37 °C for 3 hours, cells were harvested, washed with FACS buffer and stained with cell-surface markers (anti-CD8 FITC, anti-Thy1.1 PB, anti-Siglec-7 PE, and anti-Siglec-9 APC). After that, cells were fixed and permeabilized, followed by intracellular cytokine staining (anti-IFNγ PE/Cy7, anti-TNFα AF700, anti-IL2 PerCP/Cy5.5 and anti-GZMB APC/Cy7) and flow cytometry analysis.

#### In vivo Siglec-mediated P14 T cell therapy

SigE^KO^ mice were inoculated with B16-GP33 tumor cells (1×10^6^ cells per mouse as described in **Tumor models and treatments**). On day 5, mice were received with adoptive transfer of P14 T cells expressing EV or Siglec-7/9 (3×10^6^ T cells per mouse, as described above) and PBS. Tumor growth was measured every two days, and mice were humanely euthanized at the experimental endpoints as described in **Tumor models and treatments**.

#### hMDM-mediated PBMC CD8^+^ T cell activation

Anti-human CD3 (BioLegend, clone OKT3) was pre-coated in the 96-well flat-bottom plate at 100 ng/mL in 50 μL PBS per well at 4 °C overnight. hMDMs were pre-treated with PBS or 30 nM Sig7/9de for 24 hours prior to use. Autologous CD8^+^ T cells were isolated from PBMCs using EasySep human CD8+ T cell isolation kit (TEMCELL, Cat# 17953) and stained with 2 μM CFSE (BioLegend) in PBS at r.t. for 5 min, followed by washes with complete medium. 1×10^5^ CFSE-labeled CD8^+^ T cells and 1×10^4^ hMDMs in each 100 μL RPMI complete medium were mixed and seeded into OKT3 pre-coated 96-well flat-bottom plate. T cell surface expression levels of CD69, CD25 and PD-1 were measured by flow cytometry at 6, 12 and 24 hours. After 72 hours of coculture, T cell proliferation was assessed by CFSE dilution, and cytokine release including IFNγ and TNFα in the supernatant was determined by ELISA according to the manufacturer’s instructions.

#### Jurkat T cell stimulation

Jurkat T cells expressing variants, including EV, Siglec-7 and Siglec-9, were rested in RPMI compete medium overnight prior to use. 2×10^4^ cells in 200 μL complete medium were stimulated with plate-coated OKT3 (as described above) and soluble 1 μg/mL anti-CD28 (BioLegend, clone CD28.2) for 24 hours. During stimulation, indicated Siglec-7^+^ or Siglec-9^+^ Jurkat cells were treated with 30 nM Sig7/9de. After that, cells were stained with anti-CD69 APC and anti-PD-1 PE (BioLegend) for the activation marker analysis.

#### Jurkat T cell stimulation using MDA-MB-435 cancer cells

5×10^4^ rested Jurkat T cells (EV, Siglec-7 and Siglec-9) and 2.5×10^4^ MDA-MB-435 cells were seeded in 100 μL complete medium in a 96-well flat-bottom plate, in the presence of anti-HER2 BiTE (10 nM) or anti-HER2 BiTE-sialidase (10 nM) and anti-CD28 (1 μg/mL). 30 nM Sig7/9de or anti-Siglec-7 (clone 1E8) or anti-Siglec-9 (clone mAbA) blocking antibodies were treated to indicated Siglec-7^+^ or Siglec-9^+^ Jurkat cells during stimulation. Supernatants were harvested at 24 hours, and IL-2 secretion was measured by ELISA (Biolegend Cat # 431804).

#### BMDM-mediated P14 CD8^+^ T cell activation

Splenic P14 CD8^+^ T cells were isolated and stained with CFSE as described above. SigE^KO^ and Sig7/9^+^ BMDMs (with or without Siglec-7/9 pre-degradation) were pulsed with 10 pM GP33 peptide in an ultra-low attachment plate at 37 °C for 1 hour. After that, 1.5×10^5^ CFSE-labeled P14 CD8^+^ T cells and 1.5×10^4^ SigE^KO^ or Sig7/9^+^ BMDMs (with or without Siglec-7/9 degradation) in each 100 μL RPMI complete medium were mixed and seeded in 96-well flat-bottom plates. After 48 hours of coculture, T cell proliferation and cytokine release were assessed as described above.

#### In vitro P14 T cell cytotoxicity assay

B16-GP33 (GFP^+^) cells were seeded in 96-well flat-bottom plates at a density of 5×10^3^ cells per well. After 2-3 hours of incubation, P14 T cells expressing EV and Siglec-7/9 (as described above) were added at E:T ratio of 5:1. During the coculture, indicated Siglec-7/9^+^ cells were treated with 30 nM Sig7/9de. The number of viable target cells was monitored by GFP fluorescence imaging using IncuCyte (Sartorius) over 40 hours. Live cell numbers were quantified by IncuCyte software and normalized to the number of viable target cells in the B16-GP33 only group.

#### Confocal microscopic analysis of immunological synapse using Jurkat T cell and MDA-MB-435 cancer cells

2×10^5^ rested Jurkat T cells (EV, Siglec-7 and Siglec-9) and 1×10^5^ WT or pre-desialylated MDA-MB-435 cells (GFP) were seeded in 500 μL serum-free medium on a sterile #1.5 coverslide (10 mm) in a 24-well plate, in the presence of anti-HER2 BiTE (10 nM) and anti-CD28 (1 μg/mL). Pre-desialylation of MDA-MB-435 cells was performed by treatment with VC sialidase at 37 °C for 2 hours. After coculture at 37 °C for 30 min, cells were fixed with 4% PFA in PBS at r.t. for 20 min, permeabilized with 0.25% Triton X-100 in PBS at r.t. for 13 min. Cells were then washed, blocked, stained with goat anti-Siglec-7 or -9 (R&D AF1138 and AF1139, respectively), rabbit anti-SHP1 (Cell Signaling, clone C14H6), followed by staining with AF555 Dnk anti-goat (Abcam), AF647 Dnk anti-rabbit (Abcam) and AF405 Phalloidin (Invitrogen, cat# A30104). imaging was recorded and processed as described above.

#### Western blot analysis of phosphorylation of TCR signaling using Jurkat T cell and MDA-MB-435 cancer cells

Jurkat T cells (EV, Siglec-7 and Siglec-9) were rested in RPMI 1640 medium (containing 1% FBS) at 37 °C for 1.5 hours. 7.5×10^5^ rested Jurkat cells and 7.5×10^5^ MDA-MB-435 cells were precooled on ice and mixed in 100 μL serum-free medium in a 96-well U-bottom plate, in the presence of anti-HER2 BiTE (250 nM) and anti-CD28 (10 μg/mL). The coculture was initiated by centrifugation at 400g for 1 min at 4 °C, and followed by incubation 37 °C. The reactions were stopped by RIPA lysis buffer (containing protease and phosphatase inhibitor cocktails) at indicated time points. After lysis on ice for 40 min, cell lysates were cleared by centrifugation at 15,000g for 10 min at 4 °C. Equal amounts of supernatants were subjected to SDS-PAGE and transferred to a nitrocellulose membrane by semi-dry electrophoretic transfer. The membrane was blocked and probed with the following primary antibodies: anti-pCD3ζ Y142 (Abcam, Cat# ab68235), anti-CD3ζ (Biolegend, Cat # 644101), anti-pZAP70 Y319 (Cell Signaling, Cat# 2701), anti-ZAP70 (Cell Signaling, Cat# 2705), anti-pSrc Y416 for recognizing pLck Y394 (Cell Signaling, Cat# 6943), anti-Lck (Cell Signaling, Cat# 2984), anti-pLAT Y191 (Cell Signaling, Cat# 3584), anti-LAT (Cell Signaling, Cat# 45533).

For preparation of pre-desialylated MDA-MB-435 cells, trypsinized cells were incubated with VC sialidase at 37 °C for 2 hours.

For preparation of Siglec-7 or -9 predegraded Jurkat cells, Siglec-7 and -9 expressing Jurkat cells were treated with 30 nM Sig7/9de at 37 °C for 24 hours.

### Mice

Animal studies were performed under the approved protocols in accordance with the Institutional Animal Care and Use Committee (IACUC) of Scripps Research. Mice were bred and maintained in specific pathogen-free conditions in the care of the Immunology Vivarium at Scripps Research. The mouse strains used in this study included: WT C57BL/6J (B6), B6 CD45.1^+^, and B6 CD90.1^+^D^b^GP_33–41_TCR tg (P14) were purchased from The Jackson Laboratory, SigE^KO^ and Sig7/9^+^ B6 mice 18 were obtained from Dr. Ravetch at The Rockefeller University.

Experiments were conducted using the age-matched mice at 8-12 weeks old throughout the study, with both female and male subjects included in the studies. All animals were euthanized upon reaching the humane endpoints of the experiments, such as loss of body weight and signs of distress.

### Tumor models and treatments

In syngeneic mouse tumor models, 1×10^6^ B16-GMCSF, 1×10^6^ B16-GP33, 2×10^6^ B16-GP33/ B16-GMCSF (9:1), 2×10^6^ CT-2A and 4×10^5^ MT5 cells in 50 μL serum-free medium were subcutaneously (s.c.) injected into the right flank of indicated SigE^KO^ and Sig7/9^+^ mice, respectively (as described in the text). Mice were randomly assigned to the indicated groups for treatments. For the adoptive transfer experiments, P14 T cells and their variants (as described above) in 100 μL serum-free medium were intravenously (i.v.) administered. Sig7/9*de* was intratumorally (i.t.) administered at a dose of 10 μg in 20 μL PBS per mouse. Anti-CTLA4 (BioXCell, clone 9H10) was injected intraperitoneally (i.p.) at a dose of 200 μg in 200 μL PBS per mouse. Control groups were treated with PBS (i.t. or i.v.), non-degrader control (i.t.), or isotype antibody (i.p.) as indicated. Tumor growth was measured every two days using an electronic caliper, and tumor sizes were recorded as volume (mm^3^) using the formula (*length* × *width* × *width*)/2. During the tumor measurements, the investigators were not blinded to treatment assignments. Mice were humanely euthanized at the experimental endpoints when tumor size reached ≥ 1000 mm³ (B16-GMCSF, B16-GP33 and CT-2A) or 500 mm³ (MT5). Survival analysis was performed based on the designated endpoint.

For the tumor rechallenge experiment in the MT5 PDAC model, 4×10^5^ MT5 cells in 50 μL serum-free medium were s.c. injected into the opposite flanks of tumor-free mice 28 days after clearance of the initial tumors. Tumor size was monitored every two days as described above until the tumors were rejected.

For the CD8^+^ T cell depletion experiment, mice were injected i.p. with anti-CD8a (BioXCell, clone 2.43) at a dose of 150 μg per mouse on days 5 and 10 following MT5 tumor inoculation on day 0. For memory T cell-based adoptive immunotherapy, CD8^+^ T cells were isolated and enriched from the spleens and iLNs of MT5 tumor-free Sig7/9^+^ mice. Control CD8^+^ T cells were isolated and enriched from WT B6 mice at day 60 after infection with LCMV Armstrong. The CD8^+^ T cells were then i.v. administered to naive Sig7/9^+^ mice (3.5×10^6^ cells per mouse). After one day, 4×10^5^ MT5 cells were s.c. injected into the right flank of these mice.

### Tissue processing and immunophenotyping

Mouse tumors, tumor dLNs and spleens were dissociated mechanically through a 70 μm cell strainer. Samples were centrifuged at 500*g* for 5 min at 4 °C, and resuspended in new RPMI complete medium, followed by passing through a 40 μm cell strainer to afford the single-cell suspensions. RBCs were lysed in splenocytes and blood samples using ACK buffer. The resulting single-cell suspensions were used either for *ex vivo* re-stimulation or for staining with fluorescent antibodies for immunophenotyping (see below). Cells were first incubated with Fc blocker (BioLegend) in FACS buffer on ice for 10 min, then stained with fluorescent antibodies in FACS buffer on ice for 30 min, followed by staining with Ghost Dye Violet 510 (VWR) in PBS at r.t. for 10 min to preclude dead cells. Intracellular cytokine staining was performed by fixation and permeabilization after cell-surface staining. Gating strategies are shown and described in the text. The following antibodies were used in this study: CD45-PO (Invitrogen), CD45.2-PE (BioLegend), CD8α-PE (BioLegend), CD8α-PE/Cy7 (BioLegend), CD4-PerCP/Cy5.5 (BioLegend), CD4-FITC (BioLegend), NK1.1-PB (BioLegend), CD11b-APC (BioLegend), CD11b-PB (BioLegend), F4/80-PE/Cy7 (BioLegend), Ly6C-AF700 (BioLegend), Ly6C-FITC (BioLegend), Ly6G-AF700 (BioLegend), MHC-II-APC/Cy7 (BioLegend), CD11c-FITC (BioLegend), CD11c-PerCP/Cy5.5 (BioLegend), PD1-APC (BioLegend), Ly108-PE (BioLegend), CD44-AF700 (BioLegend), CD62L-APC/Cy7 (BioLegend).

1 McCord, K. A. *et al.* Dissecting the Ability of Siglecs To Antagonize Fcgamma Receptors. *ACS Cent Sci* **10**, 315-330 (2024). https://doi.org/10.1021/acscentsci.3c00969

1 Rodrigues, E. *et al.* A versatile soluble siglec scaffold for sensitive and quantitative detection of glycan ligands. *Nat Commun* **11**, 5091 (2020). https://doi.org/10.1038/s41467-020-18907-6

## Acknowledgements

This work was supported by the NIH (R35GM139643 to P.W., R35GM152118 to K.B.S., R35GM151000 to X.Z.). M.S.M is supported by a Canada Research Chair in Chemical Glycoimmunology and funding from NSERC. J. Z. is supported by the Cancer Research Institute/Irvington postdoctoral fellowship. I.A.W. is supported in part by the Hansen Chair in Structural Biology at Scripps Research. We thank Professor Jeffrey V. Ravetch for providing SigE^KO^ and Siglec-7^+^/-9^+^/SigE^KO^ mice.

## Author contributions

Conceptualization, C.W., and P.W.; Methodology, C.W., Y.H., J.Z., Q.Z., K.A.M., M.W., Y.S., D.Z., J.Y., S.C., S.H., X.Z., K.B.S., M.S.M., and P.W.; In vivo studies, C.W. and Y.H.; Investigation and analysis, C.W., Y. H., J.Z., X.Z., M.S.M., and P.W.; Writing, C.W., M.S.M., and P.W.; Review & Editing, everyone. C.W. and Y.H. contributed equally to this work.

## Declaration of interests

None.

## Extended Data Figures

**Extended Data Fig. 1.**
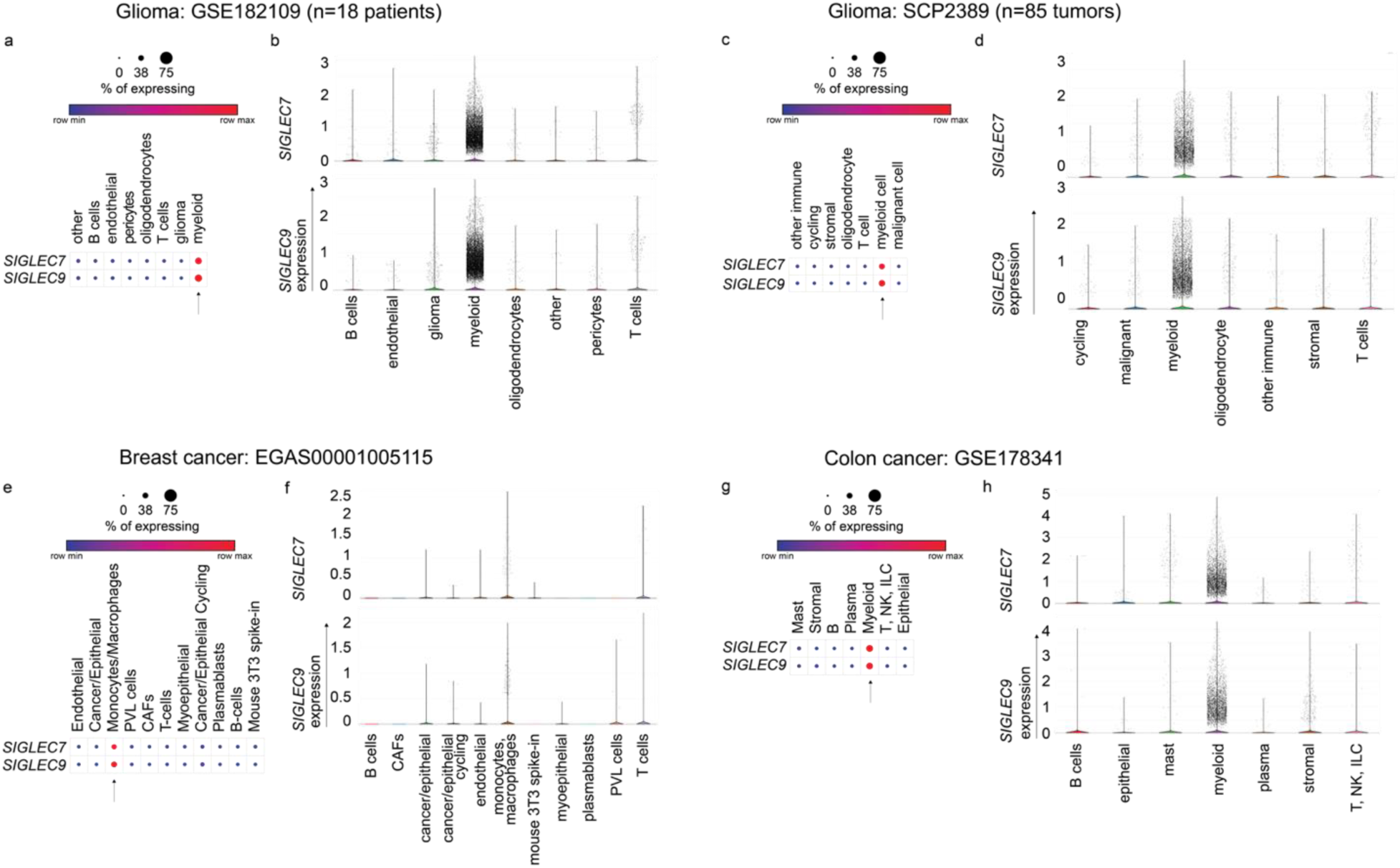
| scRNA-seq analysis of *SIGLEC7* and *SIGLEC9* expression on tumor-infiltrating cells in human cancers. Dot plots (**a**,**c**,**e**,**g**) and violin plots (**b**,**d**,**f**,**h**) presenting the expression of *SIGLEC7 and SIGLEC9* genes across annotated tumor-infiltrating cells from patient with glioma (GSE182109, SCP2389), breast cancer (EGAS00001005115) and colon cancer (GSE178341). Dot sizes represent the percentage of each cell type expressing the *SIGLEC* genes, with average gene expression values depicted on the color gradient (**a**,**c**,**e**,**g**).

**Extended Data Fig. 2.**
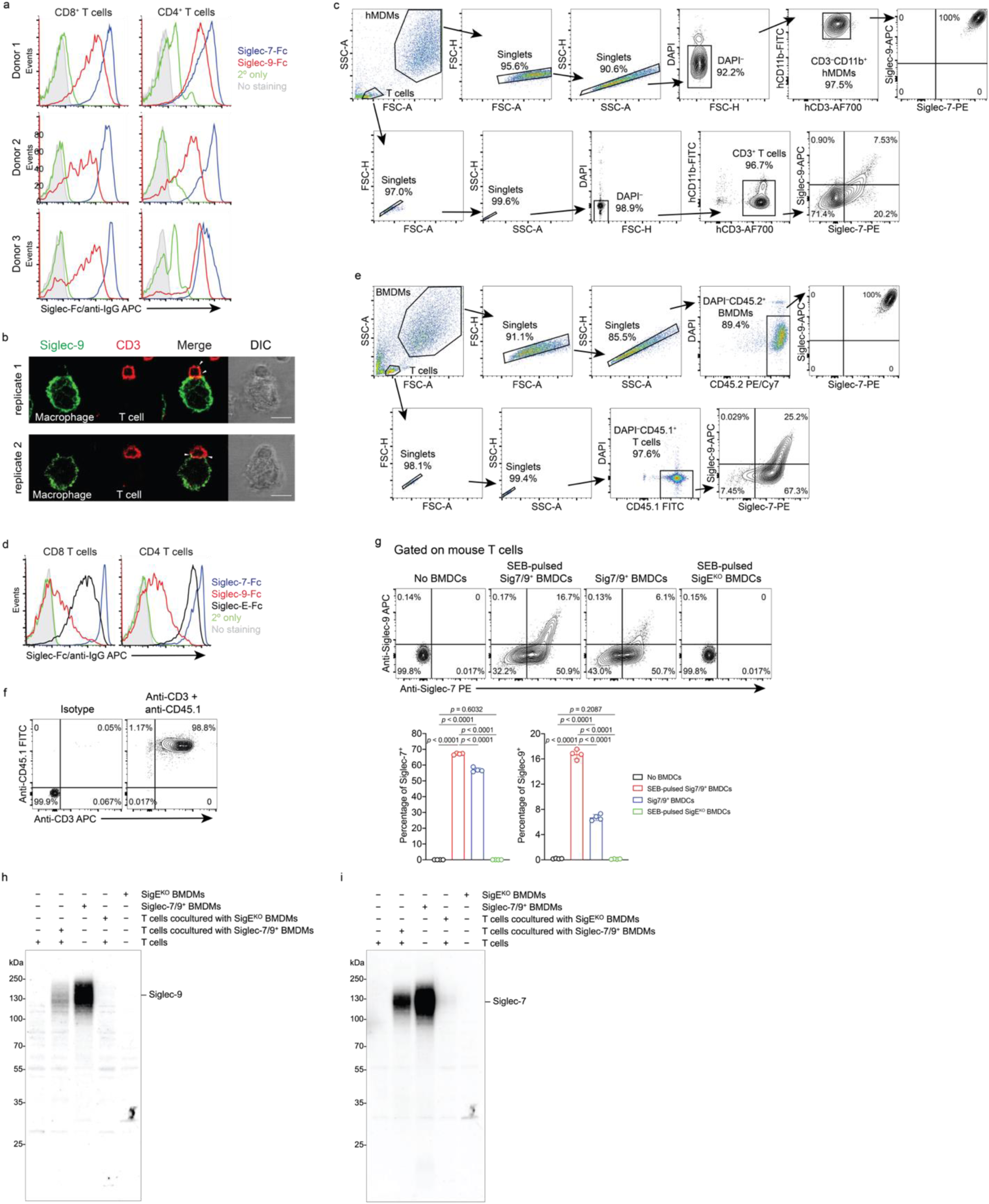
| T cells acquire Siglec-7 and -9 molecules from interacting myeloid cells via trogocytosis. **a**, Analysis of cell-surface expression of Siglec-7 and -9 ligands on CD8^+^ and CD4^+^ T cells from healthy donor PBMCs by staining with Siglec-7Fc and Siglec-9Fc, respectively. **b**, Representative fluorescence microscopy imaging analysis of Siglec-9 localization in interacting SEB-pulsed hMDMs and donor-matched PBMC T cells, which were pre-FACS sorted as Siglec-7/9^−^ population. Scale bar, 10 µm. **c**, Gating strategy for flow cytometry-based quantification of Siglec-7/-9 transfer to T cells from hMDMs, in which, the frequency of Siglec^+^ T cells was measured within the CD3^+^ T cell population after removal of debris and doublets in the absence of hMDM cells. **d**, Flow cytometry analysis of cell-surface expression of Siglec-7, Siglec-9 and Siglec-E ligands on splenic CD8^+^ and CD4^+^ T cells from WT mice by staining with Siglec-7Fc, Siglec-9Fc and Siglec-E-Fc, respectively. **e**, Gating strategy for flow cytometry-based quantification of Siglec-7/-9 transfer to WT mouse T cells (CD45.1^+/+^) from Sig7/9^+^ mouse BMDMs (CD45.2^+^), in which, the frequency of Siglec^+^ T cells was measured within the CD45.1^+^ T cell population after removal of debris and doublets in the absence of BMDM cells. **f**, Flow cytometry analysis of the purity of CD3^+^ T cells after enrichment from CD45.1^+/+^ splenic T cells. **g**, Flow cytometry analysis of Siglec-7/9 trogocytosis to WT mouse T cells after 5-min coculture with Sig7/9^+^ BMDCs *vs.* SigE^KO^ BMDCs with or without SEB pulsing. BMDC, bone marrow-derived dendritic cell. Data are mean ± s.d. Two-tailed unpaired Student’s *t*-test (**g**). **h**,**i**, Western blot analysis showing the transfer of intact Siglec-7 and -9 to WT mouse T cells from Sig7/9^+^ BMDMs compared to SigE^KO^ BMDMs.

**Extended Data Fig. 3.**
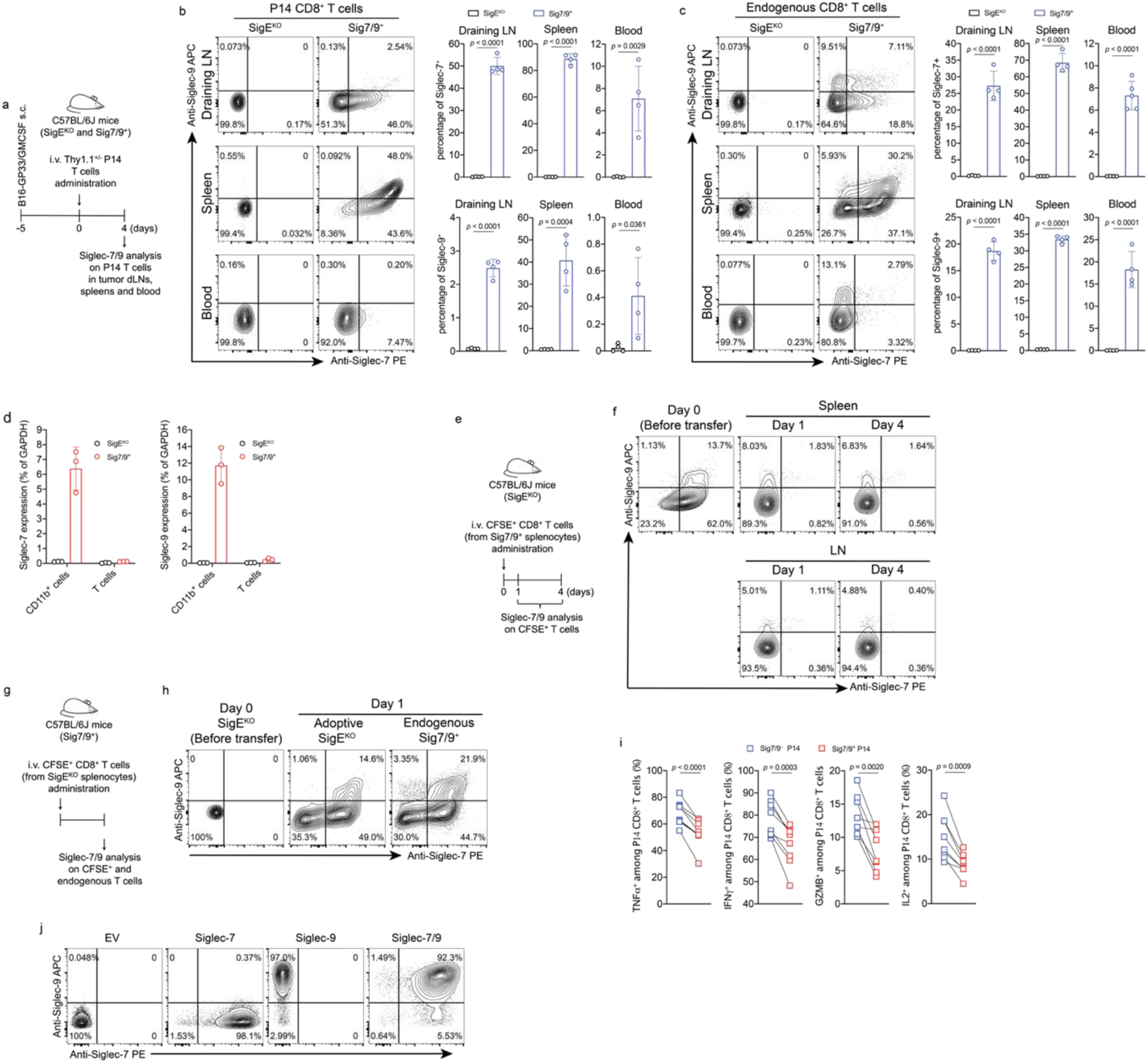
| *In vivo* Siglec-7/9 trogocytosis dampens T cell effector functions. **a**, Experimental workflow of investigation of *in vivo* trogocytosis. **b**, Analysis of Siglec-7/-9 trogocytosis by adoptively transferred P14 CD8^+^ T cells (Thy1.1) in the tumor dLNs, spleens and blood of SigE^KO^ and Sig7/9^+^ mice bearing B16-GP33/B16-GMCSF (9:1) tumors. **c**, Analysis of Siglec-7/-9 on endogenous CD8^+^ T cells. Data are mean ± s.d. Two-tailed unpaired Student’s *t*-test (**b**,**c**). **d**, Quantitative PCR analysis of Siglec-7 and -9 transcripts in isolated splenic CD11b^+^ and T cells from SigE^KO^ and Sig7/9^+^ mice. *SIGLEC* expressions were normalized based on *GAPDH* expression. **e**,**f**, Analysis of *in vivo* stability of Siglec-7/-9 on CFSE^+^CD8^+^ T cells (prepared from Sig7/9^+^ splenic cells) in the spleen and iLN tissues after adoptive transfer into SigE^KO^ recipient mice. CFSE, carboxyfluoroscein succinimidyl ester; iLN, inguinal lymph node. **g**,**h**, *In vivo* assessment of Siglec-7/-9 acquisition by SigE^KO^ T cells in spleen after adoptive transfer into Sig7/9^+^ recipient mice. **i**, Paired analysis of effector cytokine (IFNγ, TNFα, GZMB and IL2) production from Siglec-7/9^−^ and Siglec-7/9^+^ tumor dLN-infiltrating P14 CD8^+^ T cells, respectively. Paired Student’s *t*-test (**i**). **j**, Retroviral transduction of P14 T cells with individual Siglec-7 and -9, and both, respectively, characterized by staining with anti-Siglec-7 (6-434) and anti-Siglec-9 (K8).

**Extended Data Fig. 4.**
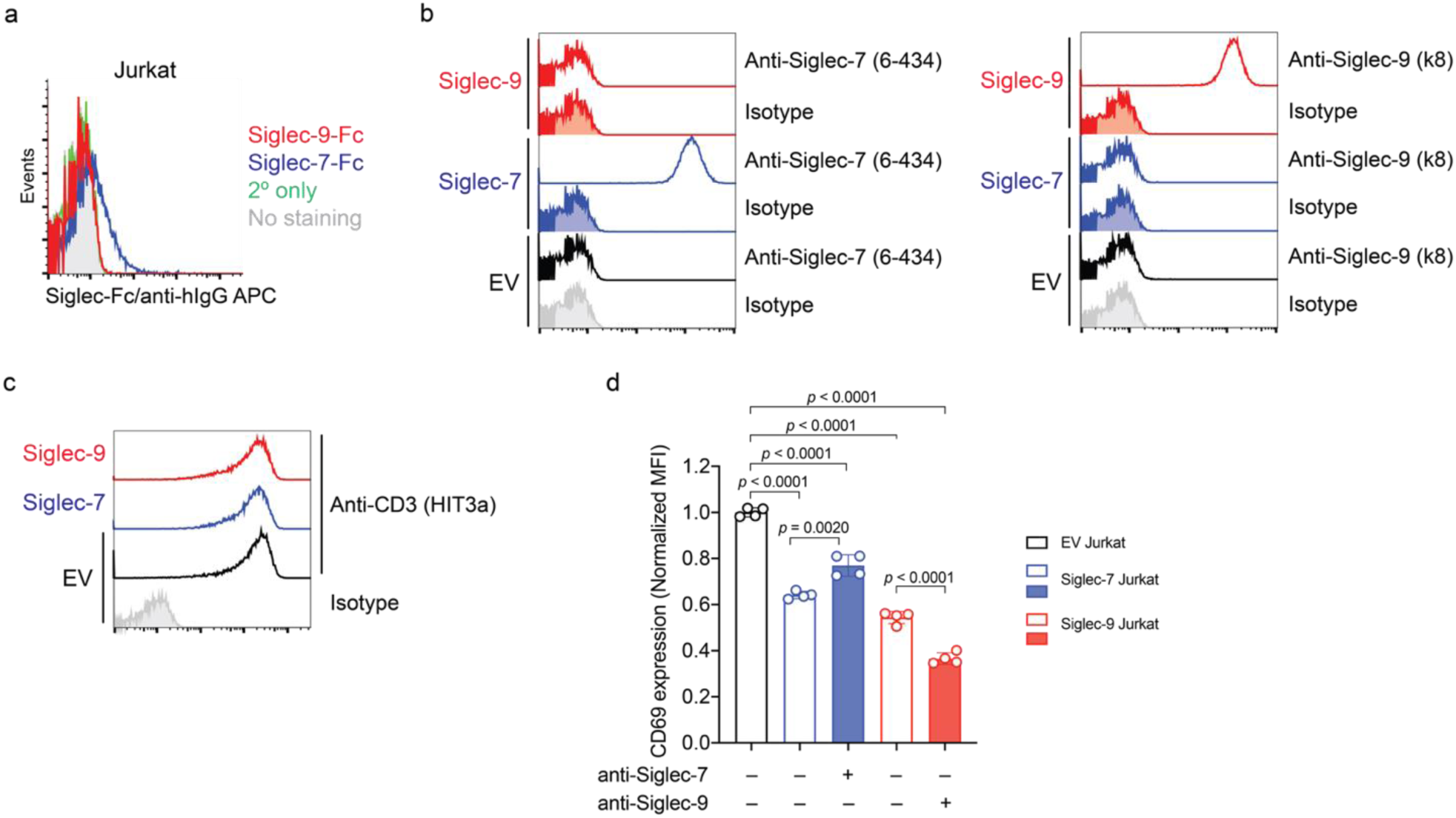
| Siglec-7 and -9 inhibit Jurkat T cell activation. **a**, Flow cytometry analysis of cell-surface expression of Siglec-7 and Siglec-9 ligands on Jurkat (E6.1) cells by staining with Siglec-7Fc and Siglec-9Fc, respectively. **b**, Lentiviral transduction of Jurkat cells with Siglec-7 WT and Siglec-9 WT, respectively, characterized by staining with anti-Siglec-7 (6-434) and anti-Siglec-9 (K8). **c**, Evaluation of CD3 expression among EV, Siglec-7 and Siglec-9 expressing Jurkat cells by staining with anti-CD3 (clone HIT3a). **d**, Assessment of Jurkat T cell (EV, Siglec-7 and Siglec-9) activation by OKT3/anti-CD28 stimulation for 24 hours, with or without anti-Siglec blocking antibody treatment (anti-Sig7 IE8 or anti-Sig9 mAbA). EV, empty vector. Data are mean ± s.d. Two-tailed unpaired Student’s *t*-test (**d**).

**Extended Data Fig. 5.**
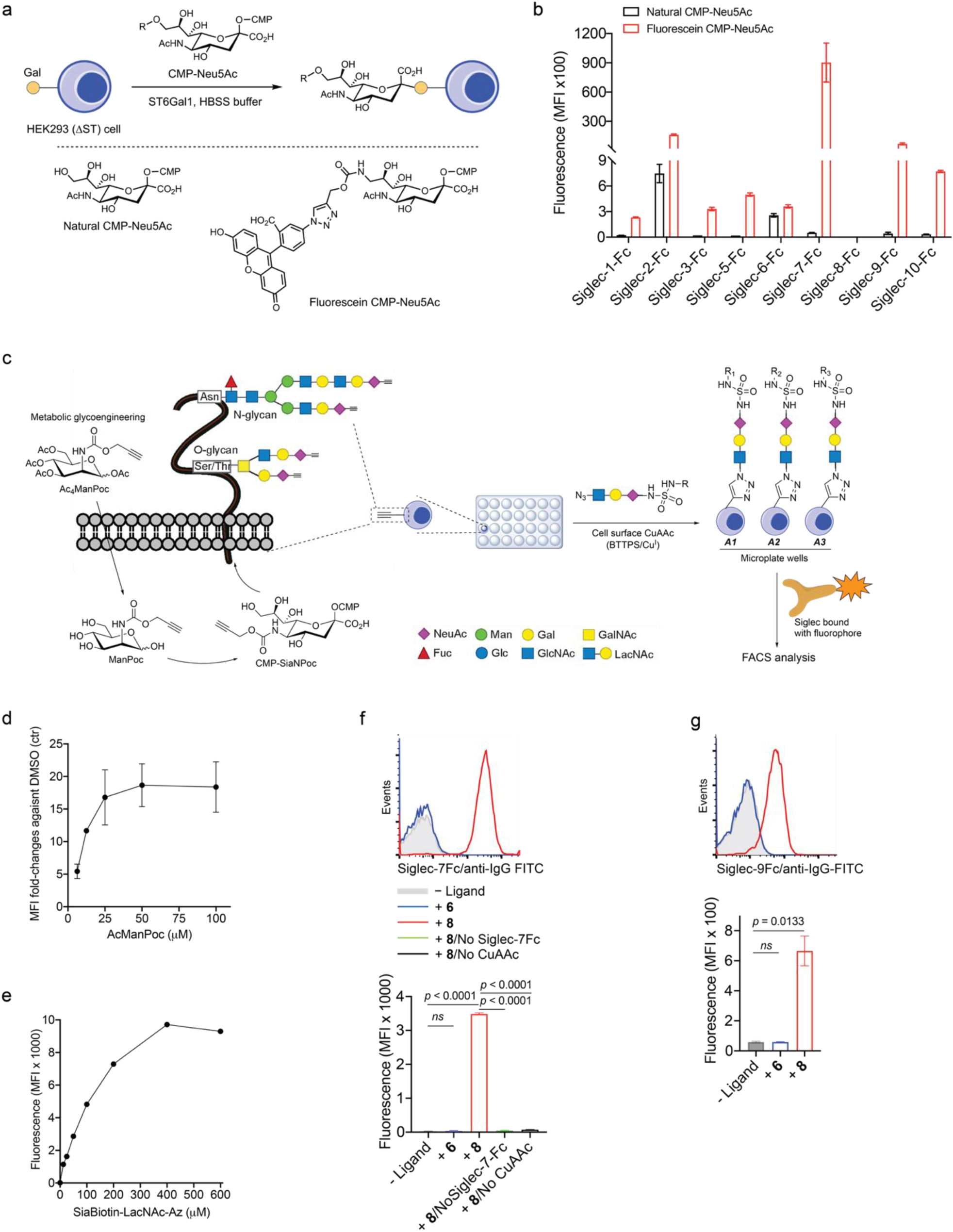
| Design and synthesis of high-affinity and selective Siglec ligands via cell-surface CuAAC assay. **a**,**b**, Evaluation of cross-binding of a reported Siglec-7 high-affinity ligand (fluorescein-modified Neu5Ac) towards a panel of human Siglecs. Fluorescein-Neu5Ac was transferred from the CMP-fluorescein-Neu5Ac donor to cell surface of sialic acid-depleted HEK293 (ΔST) cells using recombinant ST6Gal1 in HBSS buffer (**a**). Neu5Ac-installed cells were probed with fluorophore-bound Siglec-Fc chimeras for flow cytometry analysis of binding events (**b**). ST, sialyltransferase. **c**, General scheme for biocompatible cell-surface Neu5Ac ligand screening assay enabled by BTTPS-accelerated CuAAC, in which, Jurkat cells (No Siglec expression) that were metabolically labeled with Ac_4_ManPoc in complete growth media to incorporate alkynylated sialic acid onto the cell surface, were seeded to 96-well microplate in PBS buffer containing 1% FBS, pre-mixed CuSO_4_/BTTPS and Neu5Ac-LacNAc-azide ligands, followed by sodium ascorbate to initiate CuAAC to install Neu5Ac ligands onto the cell surface in a multivalent context. The modified cells were probed with fluorophore-bound Siglec-Fc chimeras for flow cytometry analysis. Man, mannose; GalNAc, *N*-acetyl-galactosamine; Fuc, fucose; Glc, glucose; Ac_4_ManPoc, *N*-propargyloxycarbamate-1,3,4,6-tetra-*O*-acetyl-manosamine; CuSO_4_, copper(II) sulfate; CuAAc, Cu(I)-catalyzed azide–alkyne cycloaddition; BTTPS, 3-[4-{(bis[(1-*tert*-butyl-1H-1,2,3-triazol-4-yl)methyl]amino)methyl}1H-1,2,3-triazol-1-yl]propyl hydrogen sulfate; FACS, fluorescence-activated cell sorting. **d**, Jurkat cells were incubated with Ac_4_ManPoc at various doses for 3 days, followed by conjugation with biotin azide via cell-surface BTTPS-accelerated CuAAc and staining with APC streptavidin for flow cytometry analysis. Biotin azide, PEG4 carboxamide-6-azidohexanyl biotin. **e**, Alkyne-labeled Jurkat cells were reacted with biotinylated Neu5Ac-LacNAc-azide at different concentrations via BTTPS-accelerated CuAAc, followed by staining with APC streptavidin for flow cytometry analysis. **f**,**g**, Histogram and MFI analysis of benzothiazole-modified Neu5Ac-LacNAc ligand (**8**) installed on Jurkat cells probed with the recombinant Siglec-7 and -9 Fc chimeras (based on the optimized conditions from **c**,**d**,**e**) in comparison with ‘No ligand’ treatment and natural Neu5Ac-LacNAc (**6**), in the presence or absence CuAAc click chemistry. The elimination of individual components in the binding assay completely negated the observed binding events, validating their authenticity (**f**). Data are mean ± s.d. Two-tailed unpaired Student’s *t*-test. *ns*, not significant (**f**,**g**).

**Extended Data Fig. 6.**
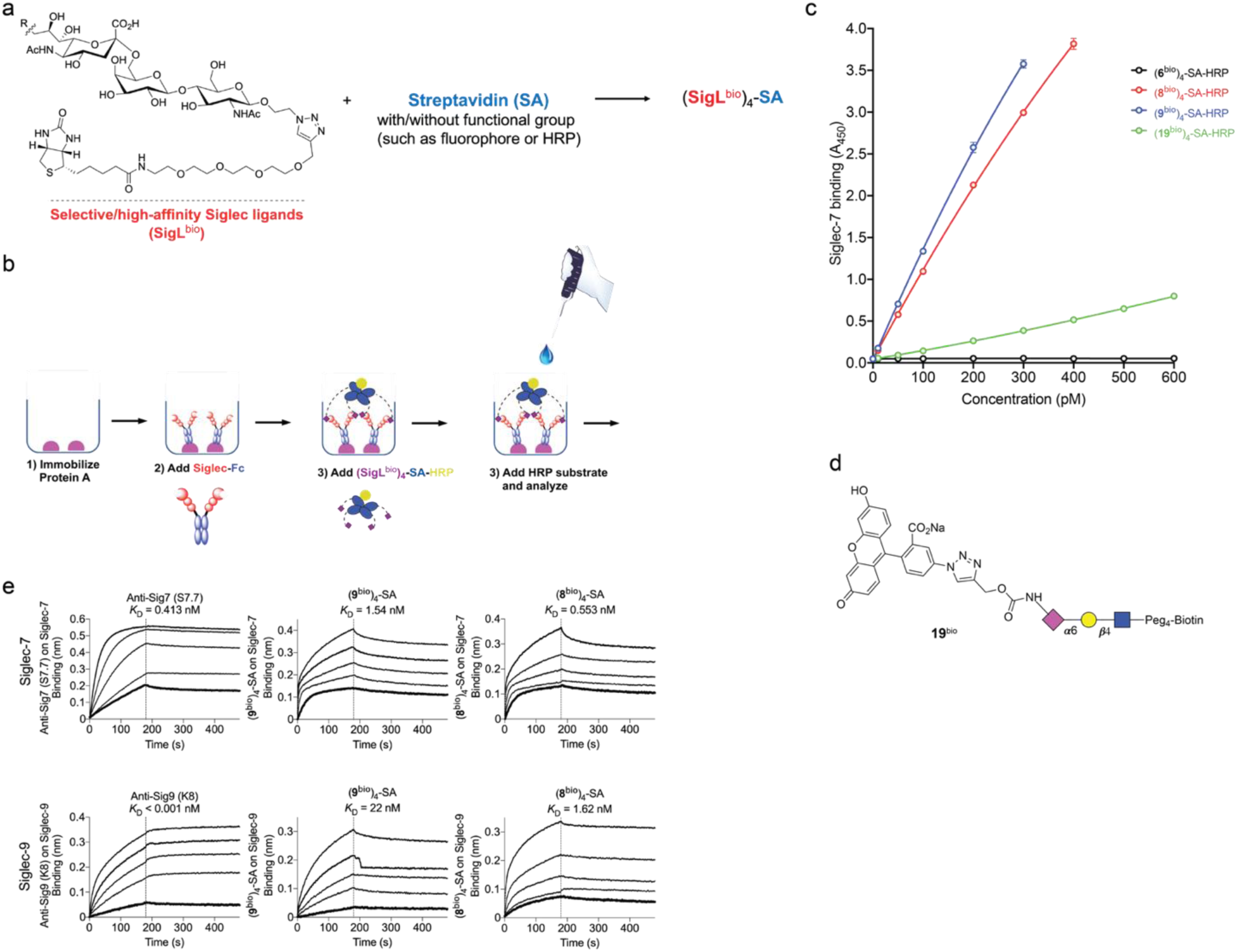
| SigL tetramers target Siglec-7/-9 comparable to antibodies. **a**, Schematic design and preparation of Siglec ligand (SigL) tetramer by combining four-equivalent biotinylated Neu5Ac-LacNAc ligands to each streptavidin (SA), in which, SA can be conjugated with a fluorophore or HRP for functional studies. **b**, The diagram showing the ELISA-like assay for measurement of SigL binding affinity, by immobilizing Siglec-Fc to protein A-coated plate, followed by incubation with HRP-conjugated (SigL^bio^)-SA tetramer and HRP substrate for signal detection. HRP, horseradish peroxidase. **c**, ELISA assay for measurement of binding affinity of ligands **8**^bio^ and **9**^bio^ to Siglec-7 in comparison with natural ligand **6**^bio^ and reported Siglec-7 ligand **19**^bio^. **d**, Chemical structure of biotinylated Siglec-7 ligand (**19**^bio^). **e**, Biolayer interferometry (BLI) assay for measurement of binding affinity of anti-Siglec-7/-9 antibodies and (**8^bio^**)_4_-SA and (**9^bio^**)_4_-SA tetramers for binding to immobilized Siglec-7 and Siglec-9 respectively. Dotted lines show association and dissociation steps.

**Extended Data Fig. 7.**
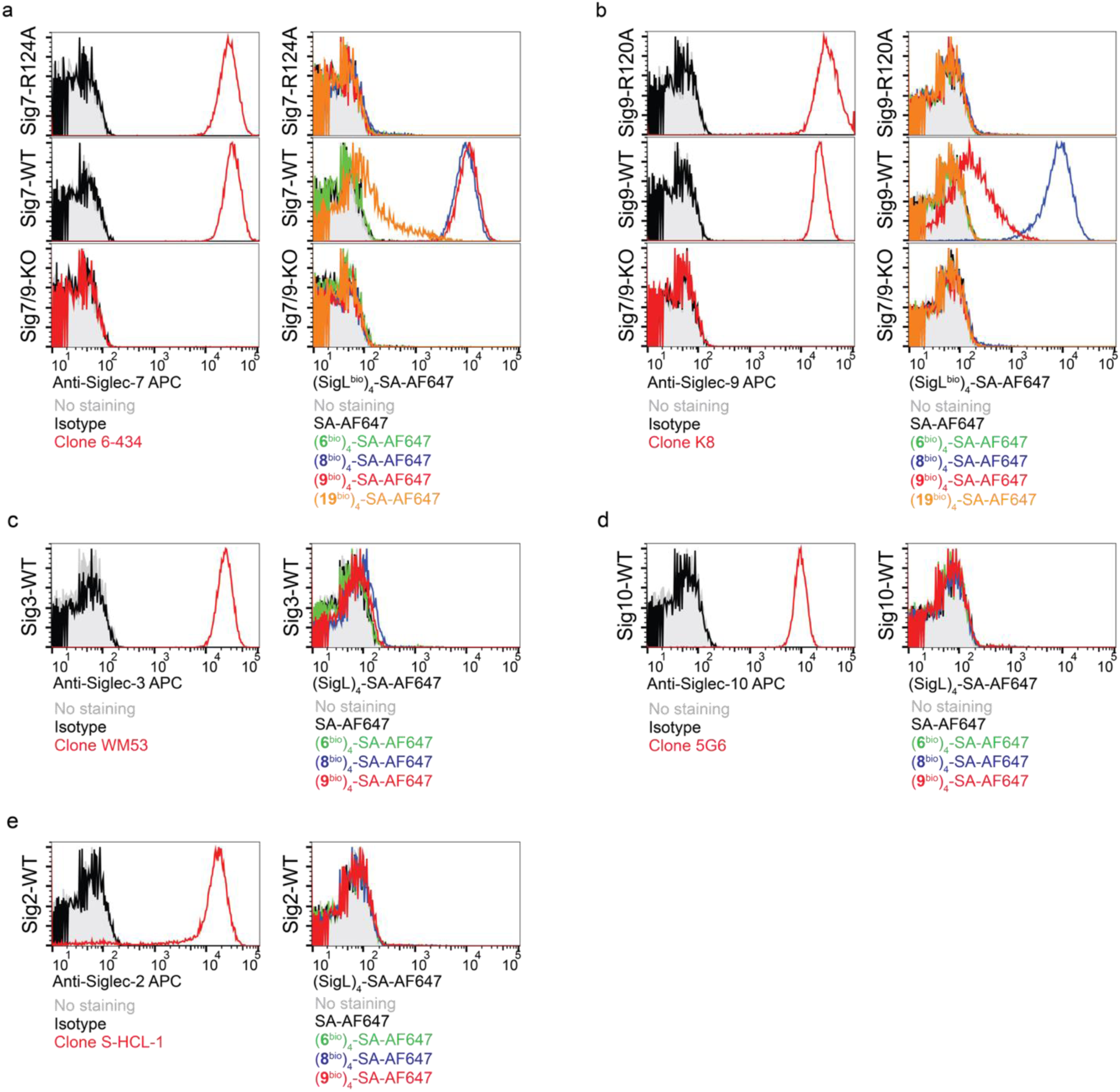
| SigL tetramers specifically label Siglec (WT)^+^ cells as an antibody surrogate. **a**,**b**, Staining of Siglec-WT (wide type), Siglec-KO (knockout) and Siglec-R mutant on U937 cells using anti-Siglec antibodies and SigL tetramers bearing AF647 dye. Data for staining Siglec-7^+^ U937 cells are shown in (**a**); Data for staining Siglec-9^+^ U937 cells are shown in (**b**). **c**, Cell surface staining of Siglec-3 WT U937 cells with anti-Siglec-3 (clone WM53) and (SigL**^bio^**)_4_-SA-AF647 tetramers. **d**, Cell surface staining of Siglec-10 WT CHO cells with anti-Siglec-10 (clone 5G6) and (SigL**^bio^**)_4_-SA-AF647 tetramers. **e**, Cell surface staining of Siglec-2 WT CHO cells with anti-Siglec-2 (clone S-HCL-1) and (SigL**^bio^**)_4_-SA-AF647 tetramers.

**Extended Data Fig. 8.**
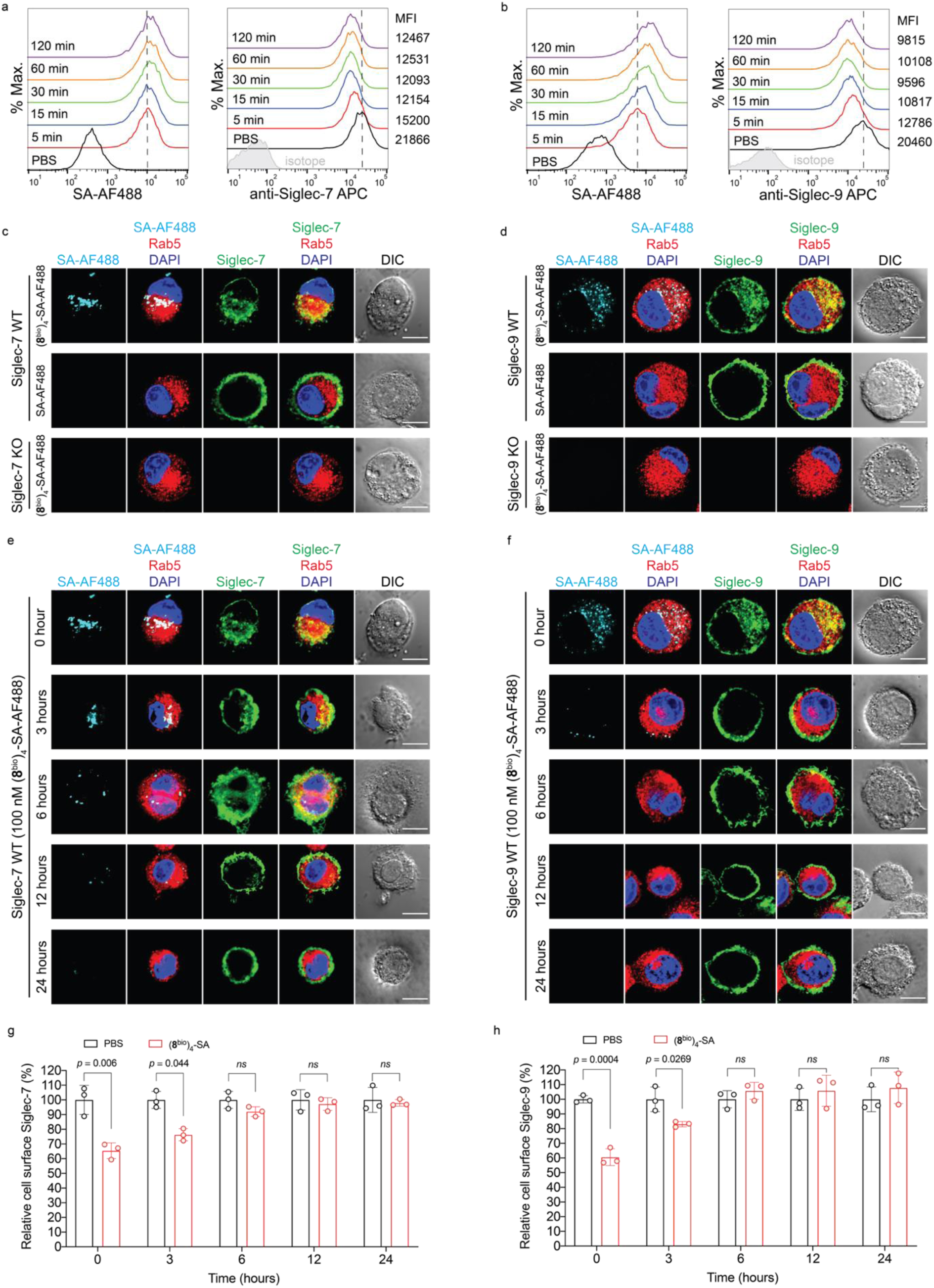
| Siglec-7 and -9 undergo rapid internalization induced by (SigL^bio^)_4_-SA tetramer and restore quickly on cell surface. **a,b**, U937-derived Siglec-7^+^ (**a**) and -9^+^ (**b**) macrophages were incubated with (**8**^bio^)_4_-SA-AF488 at 37 °C over the indicated time periods, then stained with anti-Siglec-7 and anti-Siglec-9 antibodies, followed by flow cytometry analysis. c,d, Fluorescence microscopy imaging showing that the internalized Siglec-7 (**c**)/Siglec-9 (**d**) and AF488 dye were mostly colocalized with the early endosome marker (Rab5). Scale bar, 10 µm. **e,f**, Fluorescence microscopy imaging illustrating the internalized Siglec molecules and AF488 dye in U937-derived macrophages disappear quickly following the wash away of (**8**^bio^)_4_-SA tetramer. Scale bar, 10 µm. **g,h**, Quantification of the rapid restoration of cell surface Siglec-7 (**g**) and -9 (**h**) levels on U937-derived macrophages after the removal of (**8**^bio^)_4_-SA tetramer. Data are mean ± s.d. Two-tailed unpaired Student’s *t*-test. *ns*, not significant (**g,h**).

**Extended Data Fig. 9.**
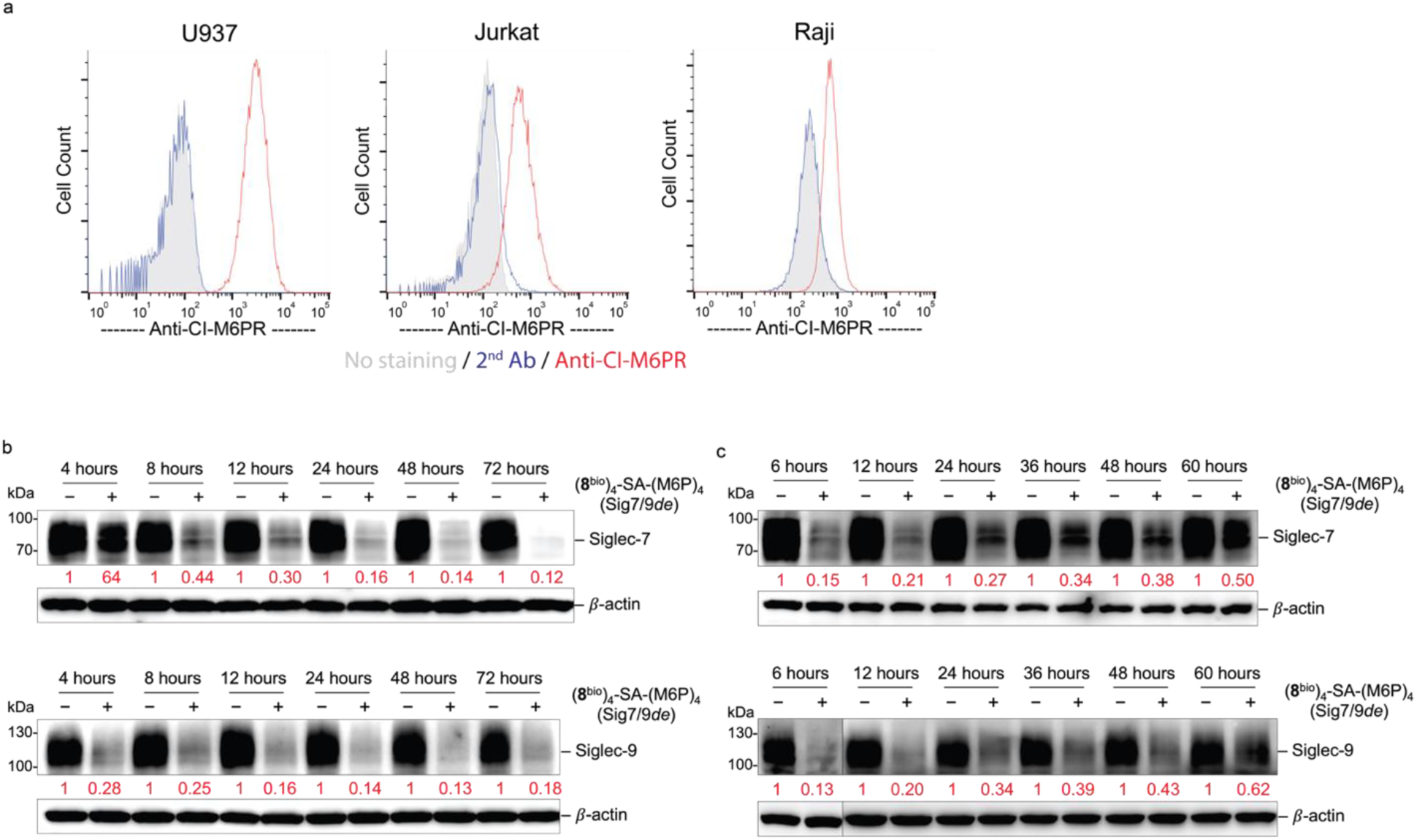
| Lysosome-targeted degradation of Siglec-7/-9 by the M6P-functionalized tetramer. **a**, Analysis of M6PR expression using anti-CI-M6PR (clone 2G11) and goat anti-mouse IgG AF488 (2^nd^ Ab). **b**, Western blot analysis of Siglec-7 and Siglec-9 proteomes in Siglec-7 WT and Siglec-9 WT U937-derived macrophages, respectively, after treatment with 30 nM (**8**^bio^)_4_-SA-M6P_4_ (Sig7/9*de*) over different time periods. **c**, Western blot analysis of recovery of Siglec-7 and Siglec-9 expression in Siglec-7 WT and Siglec-9 WT U937-derived macrophages, respectively, over different time periods, following degradation by a 24-hour treatment with 30 nM (**8**^bio^)_4_-SA-M6P_4_ (Sig7/9*de*).

**Extended Data Fig. 10.**
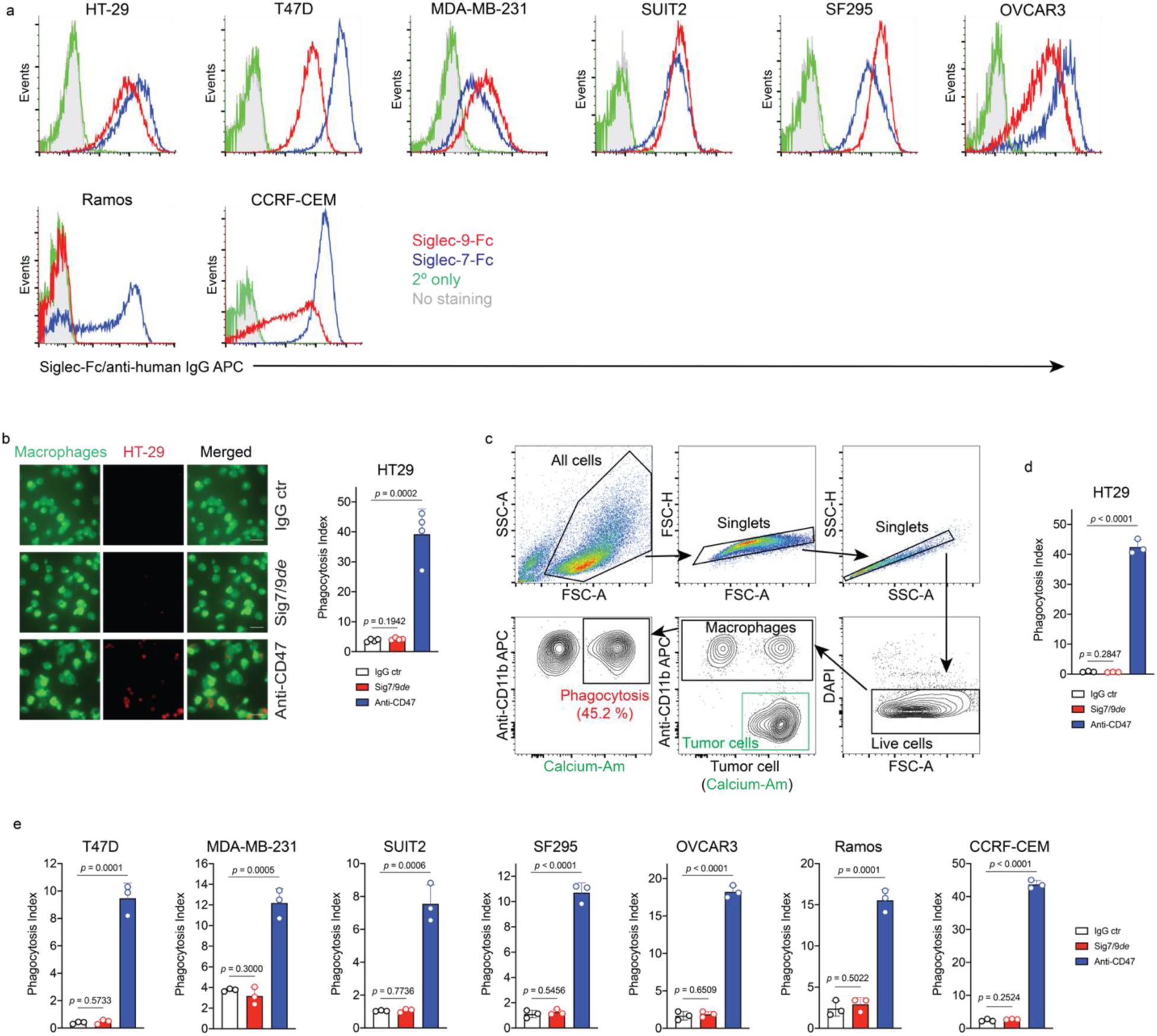
| Siglec-7/9 degradation does not improve macrophage phagocytosis of cancer cells. **a**, Flow cytometry analysis of the expression of Siglec-7 and Siglec-9 ligands on various types of tumor cell lines by staining with Siglec-7Fc and Siglec-9Fc, respectively. **b**, Fluorescence microscopy imaging-based measurement of hMDM (green) phagocytosis of pHrodo red-labeled HT29 cells in the absence or presence of Sig7/9*de* or anti-CD47 (blockade of the well-known “don’t eat me” signal CD47, which binds to signal regulatory protein-alpha (SIRPα) expressed on macrophages and dendritic cells (DCs) is used as the positive control). Scale bar, 50 µm. **c**, Gating strategy for flow cytometry-based phagocytosis assay, in which, phagocytosis was recorded by measuring the frequency of calcium-AM^+^ macrophages within the live CD11b^+^ population after removal of debris and doublets. **d**,**e**, Flow cytometry-based measurement of hMDM phagocytosis of the Siglec-7/9L^+^ cell lines of colon cancer (HT29), breast cancer (T47D and MDA-MB-231), PDAC (SUIT2), glioblastoma (SF295), ovarian cancer (OVCAR-3), B-lymphoma (Ramos) and T-ALL (CCRF-CEM) cancer cell lines, in the absence or presence of Sig7/9*de* or anti-CD47 (positive control). Data are mean ± s.d. Two-tailed unpaired Student’s *t*-test (**b**,**d**,**e**).

**Extended Data Fig. 11.**
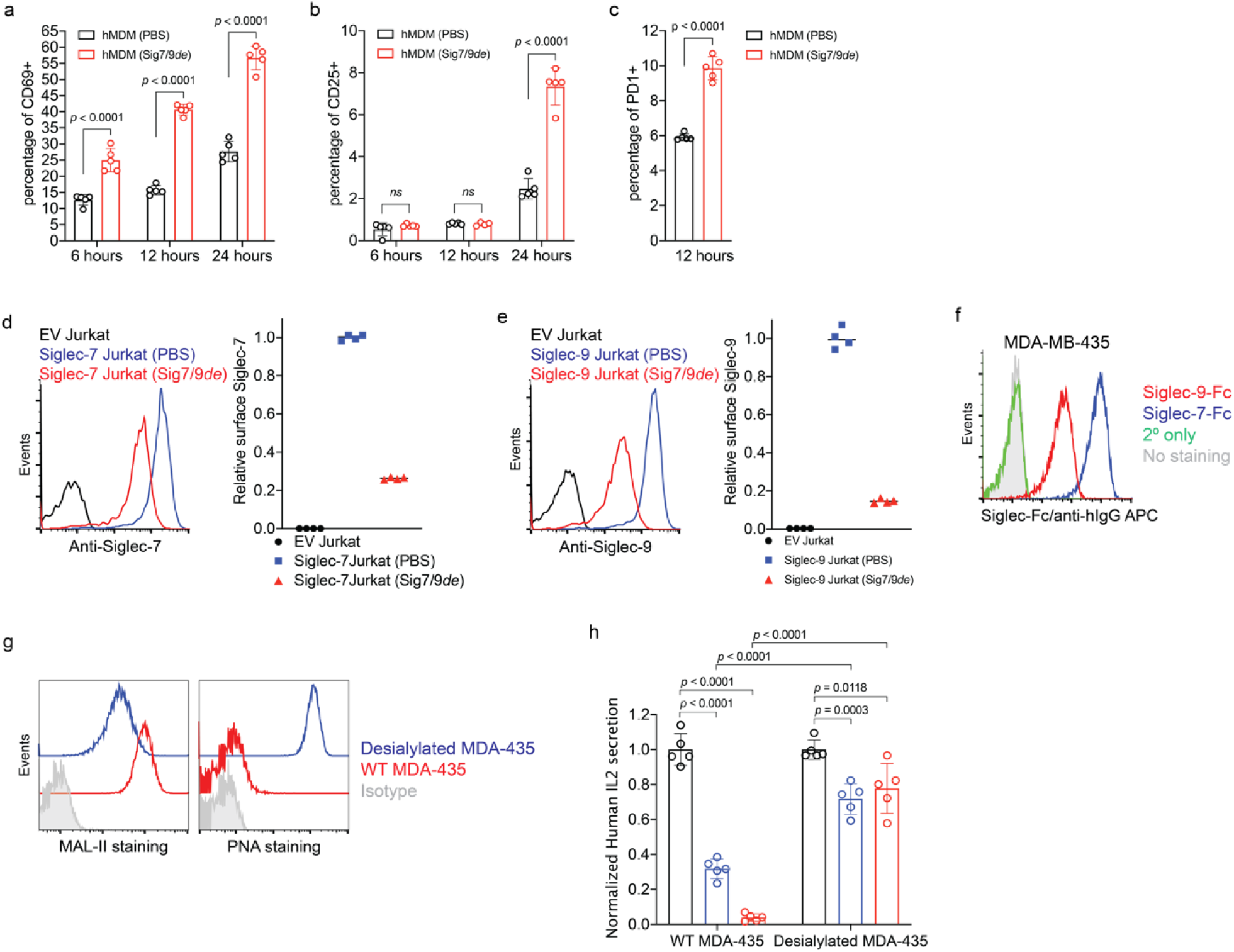
| The suppressed T cell activation induced by Siglec-7/-9 can be reversed by Siglec degradation. **a**,**b**,**c**, Assessment of activation maker (CD69 (**a**), CD25 (**b**) and PD-1(**c**)) expression on donor CD8^+^ T cells under anti-CD3 (OKT3) stimulation when cocultured with donor-matched hMDMs in the absence or presence of 30 nM Sig7/9*d*e. **d**,**e**, Analysis of cell-surface Siglec-7 (**d**) and -9 (**e**) depletion in Siglec-7 and Siglec-9 expressing Jurkat cells upon treatment with 30 nM Sig7/9*de* for 1 hour. **f**, Flow cytometry analysis of the Siglec-7 and Siglec-9 ligand expression on MDA-MB-435 cancer cells by staining with Siglec-7Fc and Siglec-9Fc, respectively. **g**, Flow cytometry analysis of sialic acid removal on MDA-MB-435 cells using anti-HER2 BiTE-sialidase by staining with MAL-II and PNA respectively. **h**, Evaluation of the cancer cell-associated *trans* ligand effect on IL2 secretion by Jurkat T cells cocultured with MDA-MB-435 cells (HER2^+^) in the presence of anti-HER2 BiTE or anti-HER2 BiTE-sialidase and anti-CD28. Data are mean ± s.d. Two-tailed unpaired Student’s *t*-test (**a**,**b**,**c**,**h**).

**Extended Data Fig. 12.**
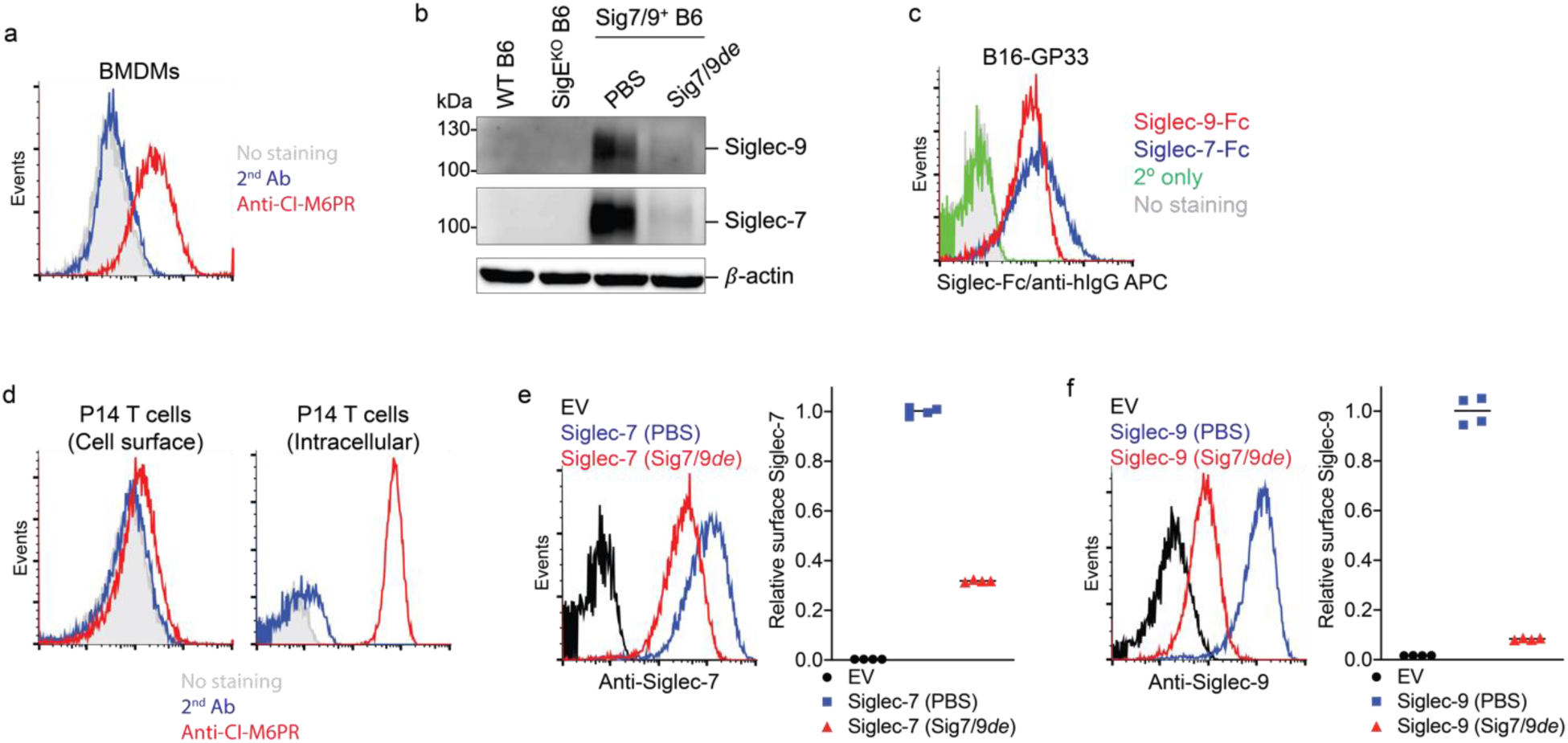
| Siglec-7/9 degradation occurs in mouse immune cells. **a**, The staining of M6PR of mouse BMDMs using anti-CI-M6PR. **b**, Western blot analysis of Siglec-7/-9 degradation in Sig7/9^+^ BMDMs in comparison to WT and SigE^KO^ controls. **c**, Flow cytometry analysis of the expression of Siglec-7 and Siglec-9 ligands on B16-GP33 cells by staining with Siglec-7Fc and Siglec-9Fc, respectively. **d**, The staining of cell surface and intracellular M6PR on P14 T cells using anti-CI-M6PR. **e**,**f**, Analysis of cell-surface Siglec-7 (**e**)/-9 (**f**) depletion in Siglec-7 and Siglec-9 expressing P14 T cells upon treatment with 30 nM Sig7/9*de* for 1 hour.

**Extended Data Fig. 13.**
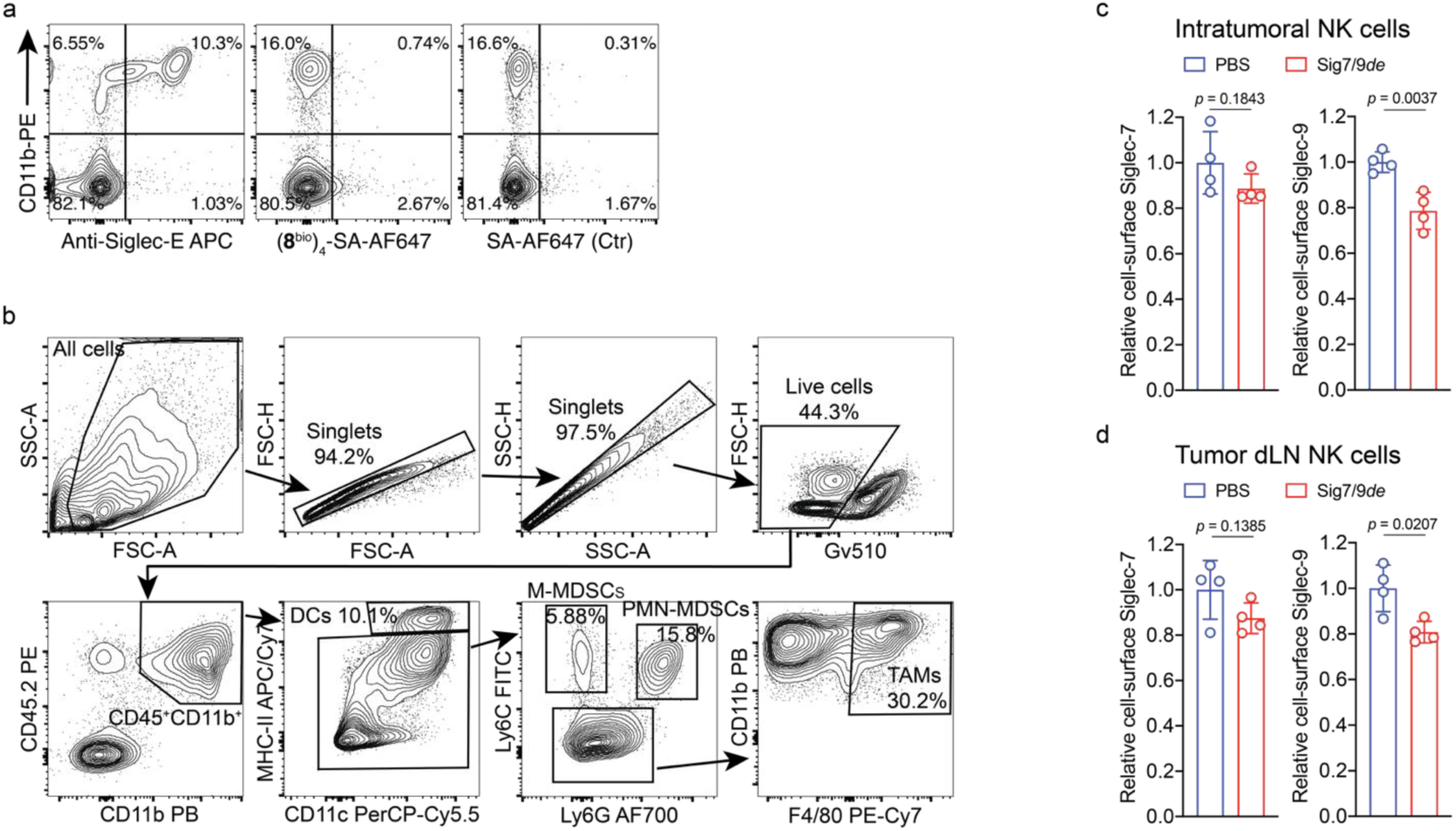
| *In vivo* degradation of Siglec-7/9 in tumor-infiltrating immune cells in Sig7/9^+^ mice. **a**, Flow cytometry analysis of Siglec-7/9 ligand (**8**^bio^) tetramer binding to peripheral blood CD11b^+^ cells from WT mice in comparison with anti-Siglec-E (clone M1304A01) staining. **b**, Gating strategy for sorting tumor-infiltrating myeloid cells in B16-GMCSF tumors inoculated in Sig7/9^+^ mice, in which, DCs, M-MDSCs, PMN-MDSCs and TAMs were characterized among CD45.2^+^CD11b^+^ population after gating out doublets and dead cells. DC, dendritic cell; M-MDSC, monocytic myeloid-derived suppressor cell; PMN-MDSC, polymorphonuclear myeloid-derived suppressor cell; TAM, tumor-associated macrophage. **c**,**d**, Assessment of *in vivo* Siglec-7/-9 depletion in tumor and tumor dLN-infiltrating NK cells following administration of 10 µg Sig7/9*de* or PBS to B16-GMCSF tumor-bearing Sig7/9^+^ mice for 2 days. NK, natural killer. Data are mean ± s.d. Two-tailed unpaired Student’s *t*-test (**c**,**d**).

**Extended Data Fig. 14.**
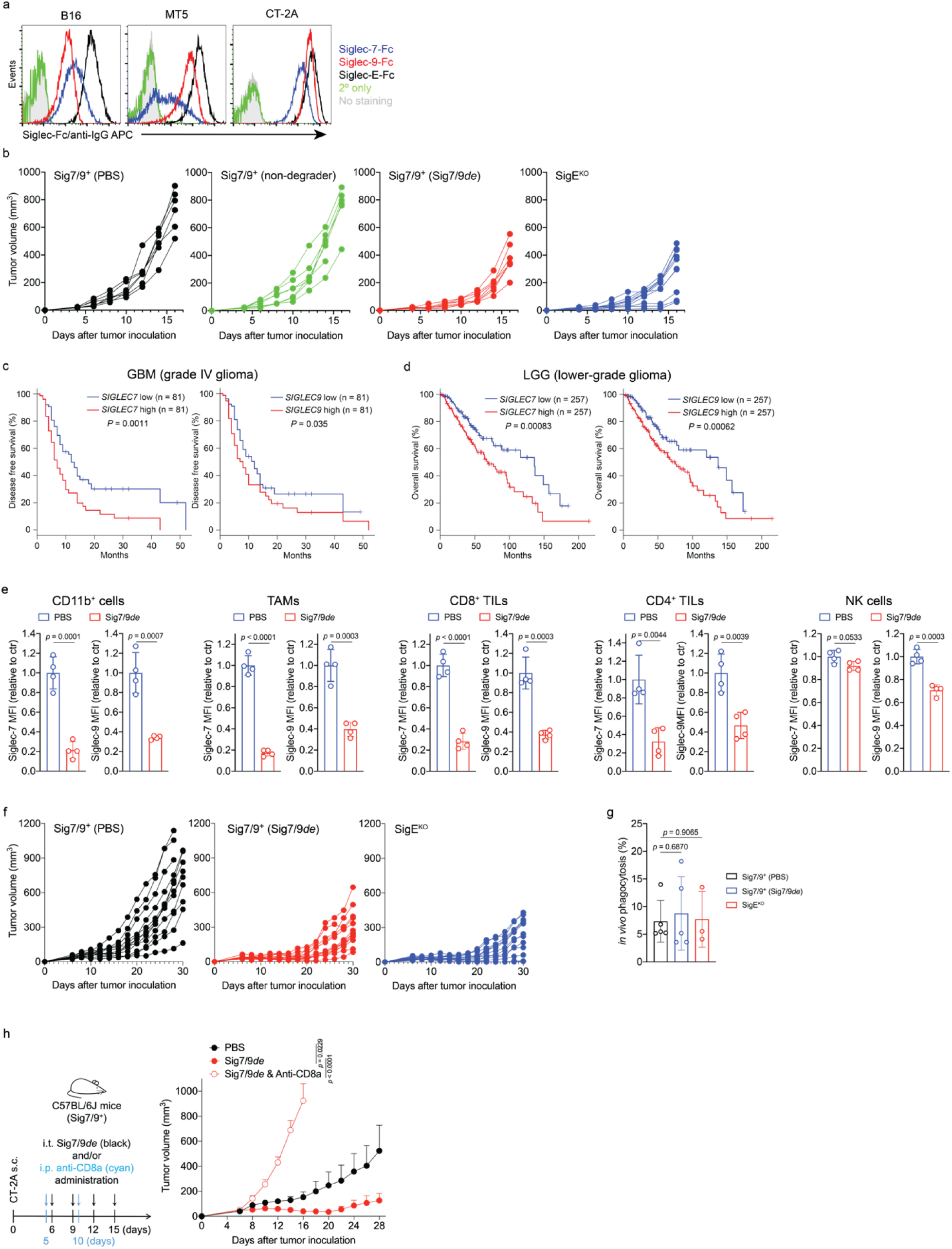
| Siglec-7/9 degradation restricts tumor growth in syngeneic mouse tumor models. **a**, Flow cytometry analysis of the expression of Siglec-7, Siglec-9 and Siglec-E ligands on three types of mouse tumor cell lines by staining with Siglec-7Fc, Siglec-9Fc and Siglec-E-Fc, respectively. **b**, Growth of individual subcutaneous B16-GMCSF tumors in SigE^KO^ (*n*= 11 mice) and Sig7/9^+^ mice treated with PBS, non-degrader ((**6**^bio^)_4_-SA-M6P_4_) and Sig7/9*de* (*n*= 7 mice per group). **c**,**d**, Relapse-free survival of patients with GBM (grade IV glioma) (**c**) and overall survival of patients with LGG (lower-grade glioma) (**d**) with high or low expression of both *SIGLEC7 and SIGLEC9* as defined by the median. The Kaplan–Meier (KM) survivals were analyzed using TCGA datasets. Two-sided *P* value computed by a log-rank (Mantel–Cox) test. GBM, glioblastoma multiforme; LGG, low-grade glioma. **e**, Assessment of Siglec-7/-9 depletion in tumor-infiltrating immune cells (including TAMs, T cells and NK cells) in CT-2A tumors in Sig7/9^+^ mice following intratumoral administration of Sig7/9*de*. **f**, Growth of individual subcutaneous CT-2A tumors inoculated in SigE^KO^ (*n*= 13 mice) and Sig7/9^+^ mice treated with PBS (*n*= 12 mice) and Sig7/9*de* (*n*= 13 mice). **g**, Assessment of *in vivo* macrophage phagocytosis of CT-2A tumor (GFP) by quantifying the percentage of GFP^+^ TAMs within tumor tissues. Data are mean ± s.d. Two-tailed unpaired Student’s *t*-test (**e**,**g**). **h**, CT-2A tumor growth in Sig7/9^+^ mice that were intratumorally administrated with PBS (*n*= 5 mice), Sig7/9*de* (*n*= 5 mice) and Sig7/9*de* in combination with anti-CD8a depleting antibody (clone 2.43) (*n*= 5 mice). Average sizes of primary tumors ± SEM are presented in cubic millimeters (mm^3^). *P* values were determined by one-way ANOVA with Dunnett’s multiple comparisons test.

**Extended Data Fig. 15.**
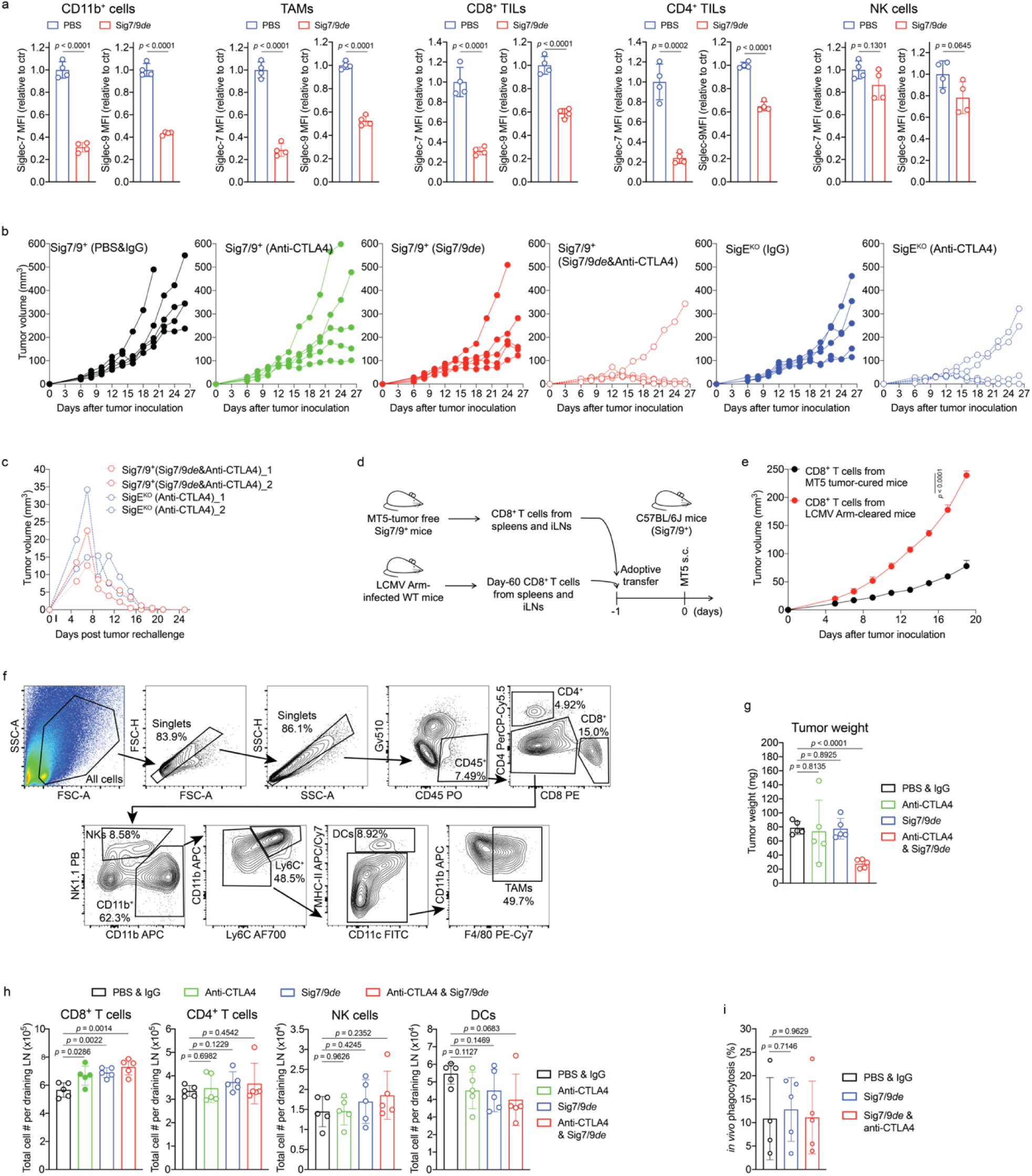
| Siglec-7/9 degradation synergizes with CTLA-4 blockade in suppressing tumor growth in a PDAC tumor model. **a**, Assessment of Siglec-7/-9 depletion in tumor-infiltrating immune cells (including TAMs, T cells and NK cells) in MT5 tumors in Sig7/9^+^ mice following intratumoral administration of Sig7/9*de*. **b**, Growth of individual subcutaneous MT5 tumors inoculated in SigE^KO^ (*n*= 5 mice per group) and Sig7/9^+^ (*n*= 5 mice per group) mice treated with PBS, Sig7/9*de*, and/or anti-CTLA-4. **c**, Rechallenge of the tumor-free mice from (**b**) with subcutaneous MT5 tumor cells in the opposing flanks (*n* = 2 in SigE^KO^ treated with anti-CTLA-4; *n* = 2 in Sig7/9^+^ co-treated with Sig7/9*de* and anti-CTLA-4). **d**,**e**, Examination of adoptive transfer of CD8^+^ T cells (isolated from MT5 tumor-cured Sig7/9^+^ mice or LCMV Arm-cleared WT mice) for the control of MT5 tumor growth in Sig7/9^+^ mice (*n*= 5 mice per group). Average sizes of primary tumors ± SEM are presented in cubic millimeters (mm^3^). **f**, Gating strategy for sorting tumor-infiltrating immune cells in MT5 tumors inoculated in Sig7/9^+^ mice treated with single PBS, anti-CTLA-4, Sig7/9*de*, and combination of Sig7/9*de* and anti-CTLA-4, in which, CD8^+^ T, CD4^+^ T, NK, F4/80^+^, Ly6C^+^ cells and DCs were characterized among live CD45^+^ population after gating out doublets and dead cells. Dendritic cell, DC. **g**, MT5 tumor weight in the MT5 tumor model in each treatment condition at day 18 (*n*= 5 mice per group). **h**, Flow cytometry analysis of numbers of tumor dLN-infiltrating immune cells in the MT5 tumor model in each treatment condition at day 18 (*n*= 5 mice per group). **i**, Assessment of *in vivo* MT5 tumor (GFP) phagocytosis by quantifying the percentage of GFP^+^ TAMs within tumor tissues from Sig7/9^+^ mice. Data are mean ± s.d. Two-tailed unpaired Student’s *t*-test (**a**,**e**,**g**,**h**,**i**).

**Extended Data Fig. 16.**
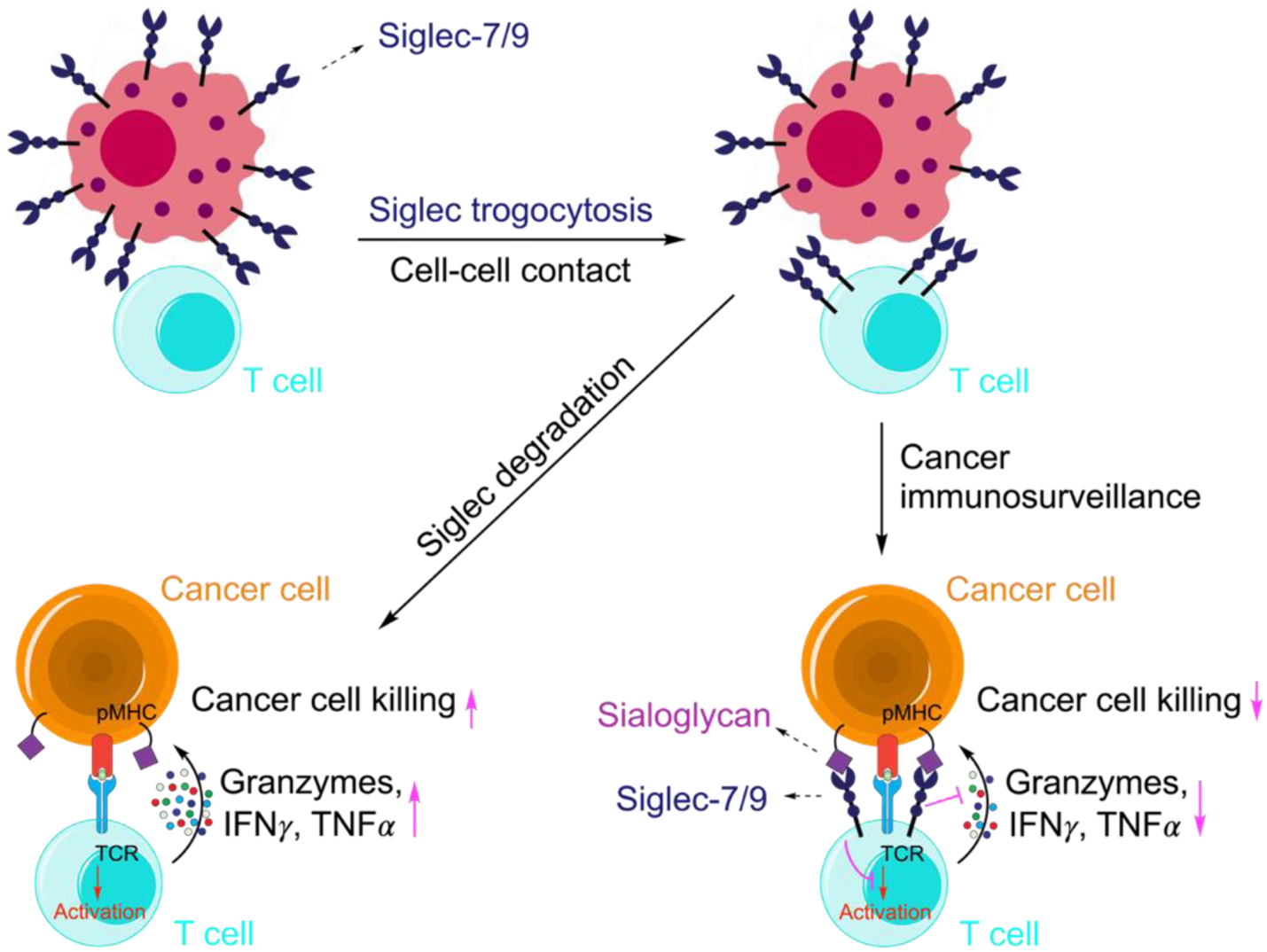
| Proposed mechanism of Siglec-7/9 trogocytosis and their inhibitory role in T cell activation. In the TME, T cells interact with neighboring Siglec-7^+^/-9^+^ myeloid cells, resulting in Siglec-7/9 trogocytosis by T cells. The acquired Siglec-7/-9 molecules suppress T cell activation, effector function, and tumor cell killing. Upon Siglec degradation, T cell effector functions are restored, allowing for better tumor control.

## Notes

### Competing Interest Statement

The authors have declared no competing interest.

